# RNA secondary structure packages evaluated and improved by high-throughput experiments

**DOI:** 10.1101/2020.05.29.124511

**Authors:** Hannah K. Wayment-Steele, Wipapat Kladwang, Alexandra I. Strom, Jeehyung Lee, Adrien Treuille, Eterna Participants, Rhiju Das

## Abstract

The computer-aided study and design of RNA molecules is increasingly prevalent across a range of disciplines, yet little is known about the accuracy of commonly used structure modeling packages in tasks sensitive to ensemble properties of RNA. Here, we demonstrate that the EternaBench dataset, a set of over 20,000 synthetic RNA constructs designed in iterative cycles on the RNA design platform Eterna, provides incisive discriminative power in evaluating current packages in ensemble-oriented structure prediction tasks. We find that CONTRAfold and RNAsoft, packages with parameters derived through statistical learning, achieve consistently higher accuracy than more widely used packages in their standard settings, which derive parameters primarily from thermodynamic experiments. Motivated by these results, we develop a multitask-learning-based model, EternaFold, which demonstrates improved performance that generalizes to diverse external datasets, including complete mRNAs and viral genomes probed in human cells and synthetic designs modeling mRNA vaccines.

## Introduction

RNA molecules perform essential roles in cells, including regulating transcription, translation, and molecular interactions, and performing catalysis.^1^ Synthetic RNA molecules are gaining increasing interest for a variety of applications, including genome editing,^2^ biosensing,^3^ and vaccination.^4^ Characterizing RNA secondary structure, the collection of base pairs present in the molecule, is typically necessary for understanding the function of natural RNA molecules and is of crucial importance for designing better synthetic molecules. Some of the most widely-used packages use a physics-based approach^5^ that assigns thermodynamic values to a set of structural features (ViennaRNA,^6^ NUPACK,^7^ and RNAstructure^8^), with parameters traditionally characterized via optical melting experiments and then generalized by expert intuition.^9^ However, a number of other approaches have also been developed that utilize statistical learning methods to derive parameters for structural features (RNAsoft,^10^ CONTRAfold,^11^ CycleFold,^12^ LearnToFold,^13^ MXfold^14^, SPOT-RNA^15^).

Secondary structure modeling packages are typically evaluated by comparing single predicted structures to secondary structures of natural RNAs^16^. While important, this practice has limitations for accurately assessing packages, including bias toward structures more abundant in the most well-studied RNAs (tRNAs, ribosomal RNA, etc.) and neglect of energetic effects from these natural RNAs’ tertiary contacts or binding partners. Furthermore, scoring on single structures fails to assess the accuracy of ensemble-averaged RNA structural observables, such as base-pairing probabilities, affinities for proteins, and ligand-dependent structural rearrangements, which are particularly relevant for the study and design of riboswitches^17, 18^, ribozymes, pre-mRNA transcripts, and therapeutics^19^ that occupy more than one structure as part of their functional cycles. Existing packages are, in theory, capable of predicting ensemble properties through so-called partition function calculations, and, in practice, are used to guide RNA ensemble-based design, despite not being validated for these applications.

High-throughput RNA structure experiments data, such as high-throughput chemical mapping^20-22^ and RNA-MaP experiments^23, 24^, offer the opportunity to make incisive tests of secondary structure models with orders-of-magnitude more constructs than previously. Unlike datasets of single secondary structures, both of these experiments provide ensemble-averaged structural properties, which allow for directly evaluating the full ensemble calculation of secondary structure algorithms, obviating the need to also evaluate the further nontrivial inference of a most-likely structure from the calculated ensemble. Furthermore, experimental data on human-designed synthetic RNA libraries has potential to mitigate effects of bias incurred in natural RNA datasets.

In this work, we evaluate the performance of commonly-used packages capable of making thermodynamic predictions in two tasks for which large datasets of synthetic RNAs have been collected via the RNA design crowdsourcing platform Eterna^25^: 1) predicting chemical reactivity data through calculating probabilities that nucleotides are unpaired, and 2) predicting relative stabilities of multiple structural states that underlie the functions of riboswitch molecules, a task that involves predicting affinities of both small molecules and proteins of interest. We find striking, consistent differences in package performance across these quantitative tasks, with the packages CONTRAfold and RNASoft performing better than packages that are in wider use. Furthermore, we develop a multitask-learning-based framework to train a thermodynamic model on these tasks concurrently with the task of single-structure prediction. The resulting multitask-trained model, called EternaFold, demonstrates increased accuracy both on held-out data from Eterna as well as completely independent datasets encompassing viral genomes and mRNAs probed with distinct methods and under distinct solution and cellular conditions.

## Results

### Evaluated packages

We evaluated commonly used secondary structure modeling packages in their ability to make thermodynamic predictions on a compilation of large datasets of diverse synthetic molecules from Eterna, which we termed EternaBench. The packages ViennaRNA (version 1.8.5, 2.4.10), NUPACK (3.2.2), RNAstructure (6.2), RNAsoft (2.0), and CONTRAfold (1.0, 2.02), were analyzed across different package versions, parameter sets, and modelling options, where available (Table S1). We also evaluated packages trained more recently through a varied set of statistical or deep learning methods (LearnToFold^13^, SPOT-RNA^15^, MXfold^14^, CycleFold^12^, and CROSS^26^), but these packages demonstrated poor performance on a subset of chemical mapping data (Figure S1), and due to their intensive runtimes, were omitted from further comparison.

### Ranking of packages based on predictions for RNA chemical mapping

Our first ensemble-based structure prediction task investigates the capability of these packages to predict chemical mapping reactivities. Chemical mapping is a widely-used readout of RNA secondary structure^20-22^ and has served as a high-throughput structural readout for experiments performed in the Eterna massive open online laboratory^25^. A nucleotide’s reactivity in a chemical mapping experiment depends on the availability of the nucleotide to be chemically modified, and hence provides an ensemble-averaged readout of the nucleotide’s deprotection from base pairing or other binding partners.^27^ We wished to investigate if current secondary structure packages differed in their ability to recapitulate information about the ensembles of misfolded states that are captured in chemical mapping experiments.

To make this comparison, we used the Eterna “Cloud Labs” for this purpose: 24 datasets of 38,846 player-designed constructs, ranging from 80-130 nucleotides in length (dataset statistics in Table S2, participant information in Table S3). These constructs were designed in iterative cycles on the Eterna platform (Figure 1A). Participants launched “projects”, each of which contained one “target structure”, and posed a design challenge or tested a hypothesis about RNA structure (project information in Table S4). The constructs designed in these labs were periodically collected and mapped using selective 2’-hydroxyl acylation analyzed by primer extension (SHAPE) and read out using the MAP-seq chemical mapping protocol.^28^ These data were returned to participants, and the results guided future lab development and construct design^29^.

**Figure 1.**
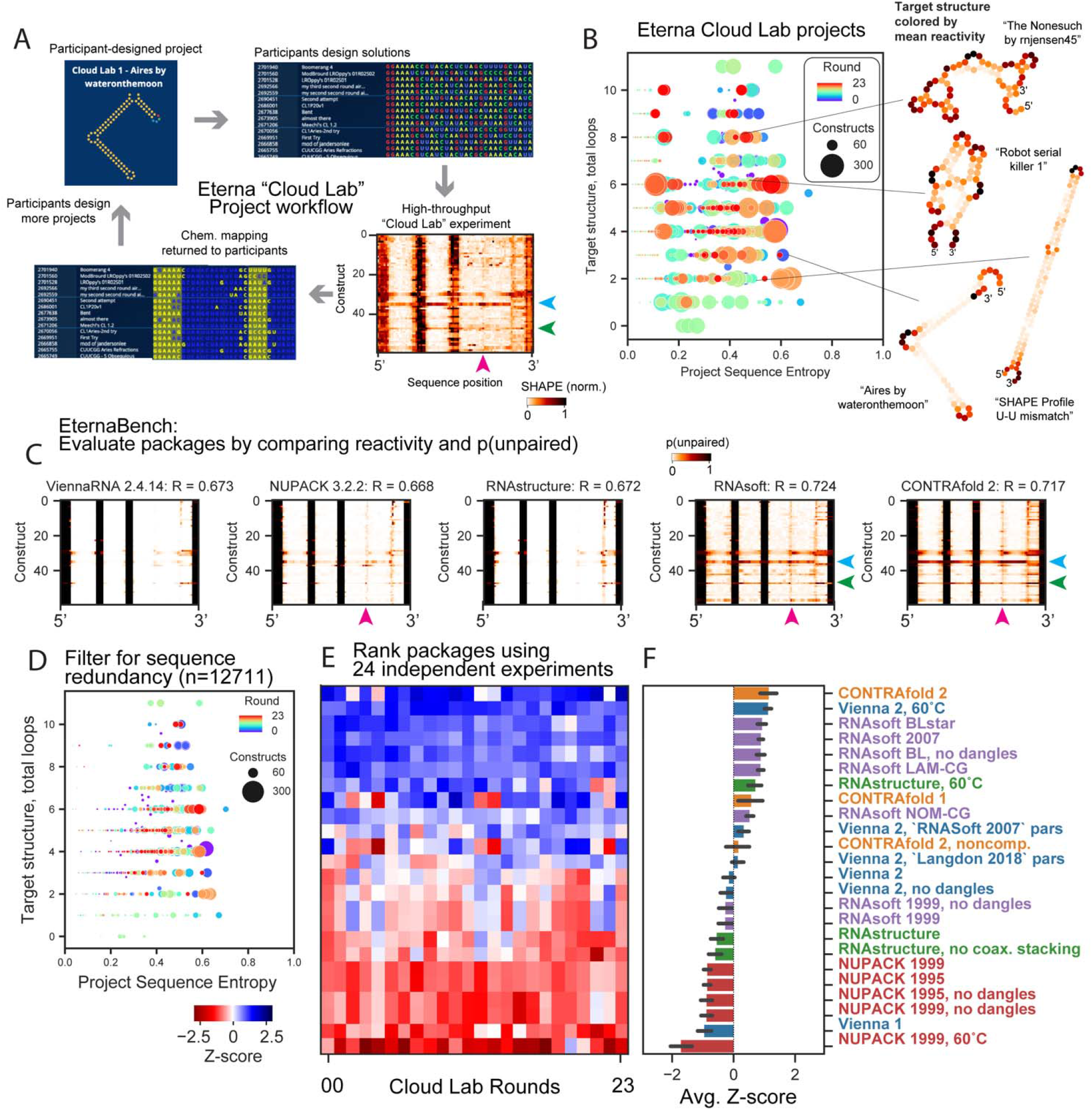
Community-science-designed RNA datasets from the Eterna “Cloud Lab” experiments identify consistent discrepancies in ensemble calculations from secondary-structure packages. (A) Workflow of cloud lab rounds: Eterna participants design “projects”, typically intended either to test RNA design challenges or energy model assumptions. Players submit solutions, all of which are synthesized in high-throughput via MAP-seq experiments. Data are returned to participants in the in-game browser, and return of results led players to design more projects. (B) Calculating the average positional entropy for all solutions collected for each project reveals that participants were able to design a diverse set of solutions, independent of target structure complexity (monitored as number of loops in the target structure). Example target structures are colored by average reactivity. (C) Example unpaired probabilities for 60 example constructs from the project “Aires” by participant wateronthemoon, for which reactivity data are shown in (A), across 5 representative secondary structure packages. Blue, green, magenta arrows indicate package predictions that recapitulate experimental partially reactive features. CONTRAfold and RNAsoft predictions for p(unpaired) have higher correlation to experimental reactivity data. (D) Analogous representation to (1B) for the EternaBench filtered dataset. (E) We compared many commonly used packages and secondary structure prediction options over 24 Cloud Lab independent experiments. We calculated the Pearson correlation coefficient and calculated the z-score across all packages evaluated for each dataset. (F) Final ranking is obtained by averaging the z-scores obtained across all datasets. Error bars represent standard deviation of z-scores across datasets.

The community of Eterna participants collectively developed highly diverse sequence libraries across target structures ranging from 0 to 12 loops (a proxy for design complexity^30^), as assessed by analyzing the positional sequence entropy of collected constructs as grouped by project (Figure 1B). Example project target structures, colored by the mean reactivity of the probed solutions, are shown in Figure 1B (inset). Some projects sought to design intricate structures, e.g., “The Nonesuch by rnjensen45” and “Robot serial killer 1”, while other participant projects focused more on better understanding experimental signals from particular structure motifs, e.g., “SHAPE Profile U-U mismatch”, which consisted of a single stem and a U-U mismatch.

Figure 1A depicts an example heatmap of SHAPE data for Eterna-player-designed synthetic RNA molecules from the project “Aires” by participant wateronthemoon. Figure 1C depicts calculated ensemble-averaged unpaired probabilities per nucleotide, *p*_*unp*_, for five example package options, plotted in the same heatmap arrangement as the experimental data in Figure 1A (see Figure S2 for heatmaps from all package options tested). In this subset of constructs, all packages are largely able to identify which regions are completely paired (*p*_*unp*_ = 0, white) or unpaired (*p*_*unp*_ = 1, black), but some packages predict *p*_*unp*_ values between 0 and 1 that more accurately reflect intermediate reactivity levels. Arrows (blue, green, magenta) indicate intermediate reactivity values that are captured by predictions from CONTRAfold and RNAsoft but not ViennaRNA, NUPACK, and RNAstructure. We quantified similarity between reactivity and p(unpaired) by calculating the Pearson correlation coefficient between the experimental reactivity values and p(unpaired) values (see Methods). In example, predictions from CONTRAfold 2 and RNAsoft BLstar for Cloud Lab Round 1 (1088 constructs) demonstrate improved correlation of R = 0.718(2), 0.724(3) (respectively) over Vienna 2, RNAstructure, and NUPACK (0.673(2), 0.671(2), 0.667(2), respectively) (Table S5). Noting that some projects had low sequence diversity, we filtered constructs to remove highly similar sequences (see Methods, Figure S3). Clustering the resulting sequences per project (Figure 1D) demonstrates that low-entropy projects were reduced in size. The final 24 EternaBench-CM datasets comprised 12,711 individual constructs.

We observed that CONTRAfold and RNAsoft generally predict that the constructs studied are more melted than the other packages predict at their default temperatures of 37 °C, even though the actual chemical mapping experiments were carried out at lower temperature (24 °C; see Methods). Motivated by this observation, we wished to ascertain if a simple change in temperature might account for differences in performance between packages. Because ViennaRNA, NUPACK, and RNAstructure packages include parameters for both enthalpy and entropy, we calculated correlations across predictions from a range of temperatures (Figure S4). We found that increasing the temperature from the default value of 37 °C used in these packages to 60 °C improved the correlation to experimental data for ViennaRNA (R=0.708(2)) and RNAstructure (R=0.707(2)), but not NUPACK (R=0.639(2)). We hence included each of these packages also at 60°C as options to test.

We established a ranking of all package options for each dataset (Figure 1E, Table S5, representative heatmaps for all datasets in Figure S5) by computing the Z-score for each package correlation in comparison to all packages tested, and averaging over all datasets (Figure 1F). The top 3 package options were CONTRAfold 2, ViennaRNA at 60°C, and RNAsoft with “BLstar” parameters. Ranking all packages with a Spearman rank correlation coefficient resulted in a similar global overall ranking (Figure S6). Overall package performance and the resulting ranking was not strongly dependent on GC content, sequence length, or total number of loops in the project target structure, which was investigated by calculating correlations and rankings when grouping constructs by project (see Methods, Figure S7).

### Ranking of packages based on predictions for riboswitch energetics

Our second ensemble-based structure prediction task involved predicting the relative populations of states occupied by riboswitch molecules. Riboswitches are RNA molecules that alter their structure upon binding of an input ligand, which effects an output action such as regulating transcription, translation, splicing, or the binding of a reporter molecule.^18, 31, 32^ We compared these packages in their ability to predict the relative binding affinity of synthetic riboswitches to their output reporter, fluorescently-tagged MS2 viral coat protein in the absence of input ligand, 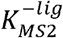 (see Methods). As with the chemical mapping datasets, each riboswitch dataset was filtered to exclude highly similar sequences (Figure S2, Table S6). These riboswitches came from two sources: the first consisted of 4,849 riboswitches (after filtering) designed by citizen scientists on Eterna^33^. The second consisted of 2,509 riboswitches (after filtering) designed fully computationally using the RiboLogic package,^34^ probed concomitantly with Eterna riboswitches.

Figure 2A depicts experimental values for 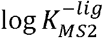 for FMN riboswitches from the RiboLogic dataset vs. predicted 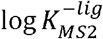 values. Again, CONTRAfold and RNAsoft BLstar packages exhibit higher correlations to the experimental data (Pearson R = 0.50(2) and 0.51(2), respectively) than ViennaRNA, NUPACK, and RNAstructure (R=0.37(2), 0.34(2), 0.36(2), respectively). Example predictions for all package options tested are in Figure S8. We evaluated performance across 12 independent experimental datasets (Figure 2B, Table S7, representative predictions in Figure S9), and obtained a ranking (Figure 2C) similar to the ranking obtained from chemical mapping data. CONTRAfold 2, RNAsoft (model “BL, no dangles”, equivalent to BLstar but without dangles), and RNAstructure 60°C were ranked as the top 3 out of the package options tested. The top ranking of CONTRAfold 2 matches the entirely independent ranking based on chemical mapping measurements of distinct RNA sequences described in the previous section. Predicting MS2 binding affinity in the presence of the riboswitch input ligand, 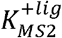, as well as the activation ratio, AR, requires computing constrained partition functions, a capability limited to Vienna RNAfold, RNAstructure, and CONTRAfold. Rankings for predicting 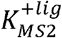 and AR followed the same trends (Figure S10, see Methods).

**Figure 2.**
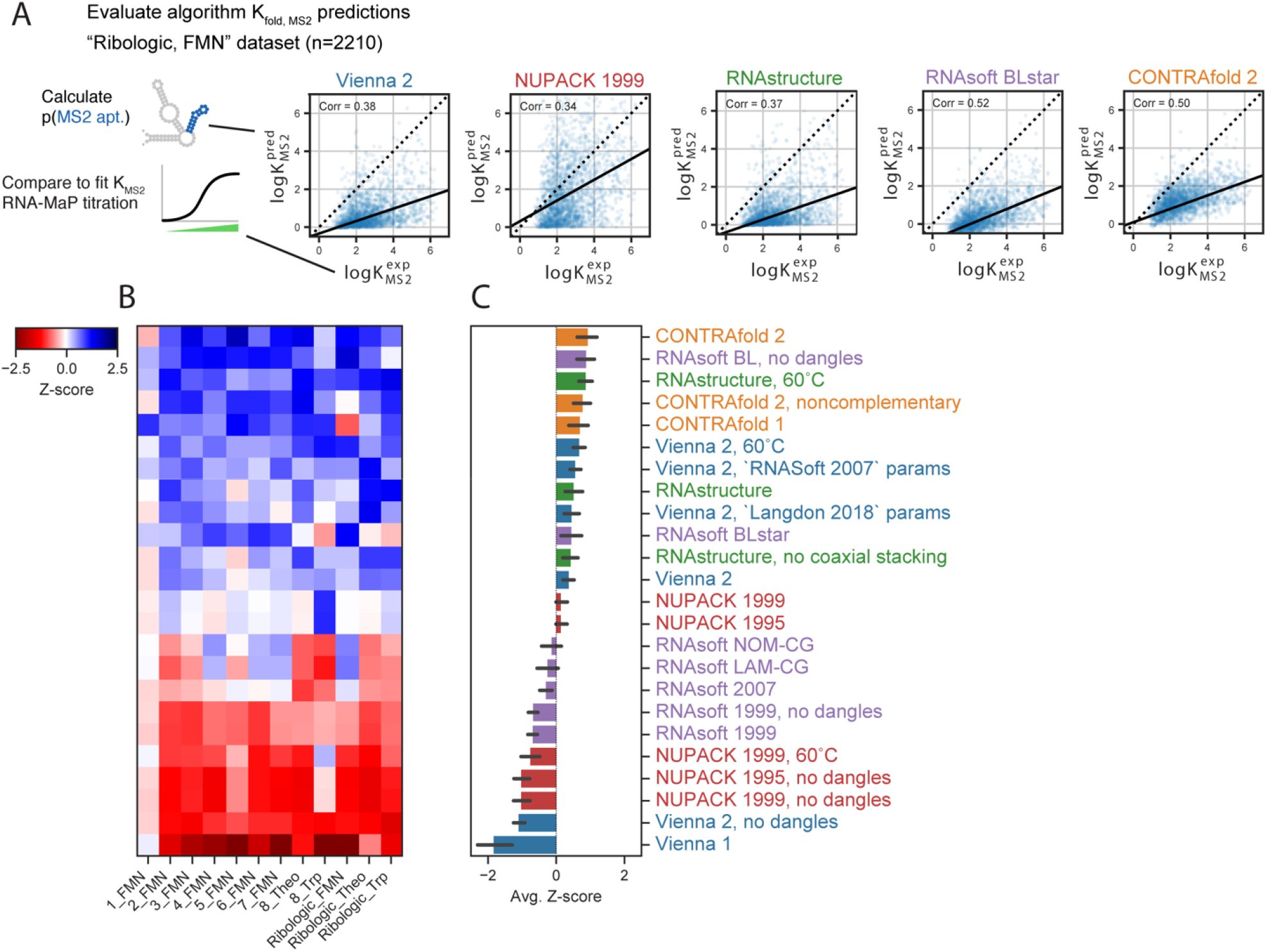
Riboswitch protein-binding predictions reveal similar package ranking. (A) Representative scatterplots for the Ribologic-FMN dataset of experimental vs. predicted values of 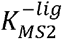. (B) Calculating the Z-scores of Pearson correlation coefficients across 12 independent datasets of riboswitches result in an overall ranking (C) consistent with the Chemical Mapping dataset.

### EternaFold gives best-of-class performance in multiple structure prediction tasks

We hypothesized that performance in both secondary structure prediction tasks above might be improved by incorporating these tasks in the process of training a secondary structure package. The RNAsoft^10, 35^ and CONTRAfold^11^ packages both take advantage of the property that the gradient of any parameter is expected counts of that feature in the ensemble, which can be readily computed in dynamic programming scheme. We generalized this framework beyond maximizing the likelihood of one single structure to matching the experimentally determined probability of a particular structural motif in the ensemble through minimizing the RMSE to riboswitch affinities for MS2 protein (see Methods). We used the CONTRAfold code as a framework to explore multi-task learning on RNA structural data, since it has previously been extended to train on chemical mapping data to maximize the expected likelihood of chemical mapping data.^36^

We trained models with a variety of combinations of data types to explore interactions in multitask training (Figure 3A), using a holdout set from the training set to determine hyperparameter weights (see Methods), and evaluated performance on separate test sets for single-structure prediction accuracy^37^, chemical mapping prediction accuracy, and riboswitch fold change prediction. To ensure a rigorous separation of training and test data, test sets for chemical mapping and riboswitch data came from completely different experimental rounds than those used in training (see Methods).

**Figure 3.**
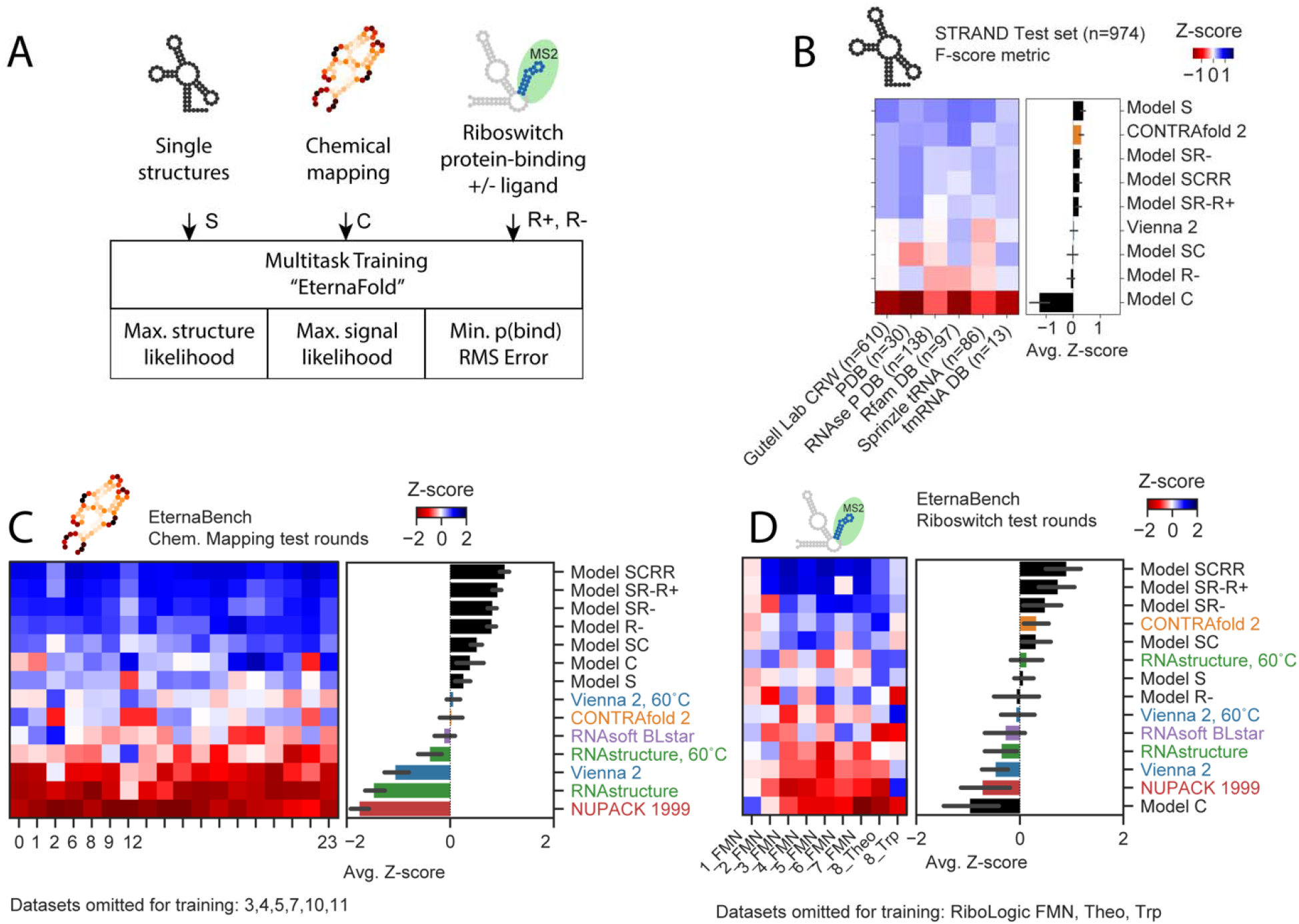
Multitask training using EternaBench datasets results in improved thermodynamic prediction. A) Scheme of data types used in multitask training and loss function used for each. B) Secondary structure prediction on STRAND test set. C) Z-score ranking over 18 test datasets for EternaBench Chemical Mapping. D) Z-score ranking over 9 Riboswitch test sets for riboswitch K_MS2_ prediction.

Comparing performance across models trained with different types of input data indicates some tradeoffs in performance. “Model S”, trained only on single-structure prediction training data, exhibited the highest accuracy on the separate single-structure prediction test set, outperforming CONTRAfold 2 and ViennaRNA 2 (Figure 3B, F-scores of 0.71(1), 0.68(1), and 0.65(1), respectively). Incorporating other data types in model training resulted in F-scores slightly worse than Model S on the single-structure prediction test set but still higher than or within error of CONTRAfold 2 (Figure 3B). Model “SCRR”, trained on four data types (single-structure data, chemical mapping, riboswitch 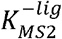 and 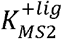) exhibited the highest performance on separate test sets for chemical mapping (Figure 3C) and riboswitch 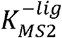 prediction (Figure 3D, data for all test sets in Table S8). We termed this SCRR model “EternaFold”.

### Independent tests confirm EternaFold performance

We wished to test if EternaFold’s improvements in correlating p(unpaired) values to chemical mapping and protein-binding data generalized to improvement in predictions for datasets from other groups, experimental protocols, and RNA molecules. We compiled 36 datasets of chemical mapping data for molecules including viral genomes^38-48^ in cells and in virions, ribosomal RNAs^43, 49, 50^ both in cells and extracted from cells, synthetic mRNAs and RNA fragments designed to improve protein expression and in vitro stability^19, 51^, and mRNAs probed in various subcellular compartments and extracted from HEK 293 cells^52^ (Figure 4A, Table S9). These datasets spanned structure probing methods different from those used in the Eterna Cloud Labs (SHAPE-CE, SHAPE-MaP, DMS-MaP-seq vs. MAP-seq) as well as a variety of chemical modifications (DMS, icSHAPE, NAI). Most of these test molecules were much longer (thousands of nucleotides) than the 85-nucleotide RNAs used as the primary training data for EternaFold. Notably, 6 of these involved the SARS-CoV-2 genome^45-47^, which came into prevalence after the development of the EternaFold model, and represented a test of novel data. Results here are from windowed folding of length 900, which resulted in the highest correlations; see Methods for evaluation of other folding models.

**Figure 4.**
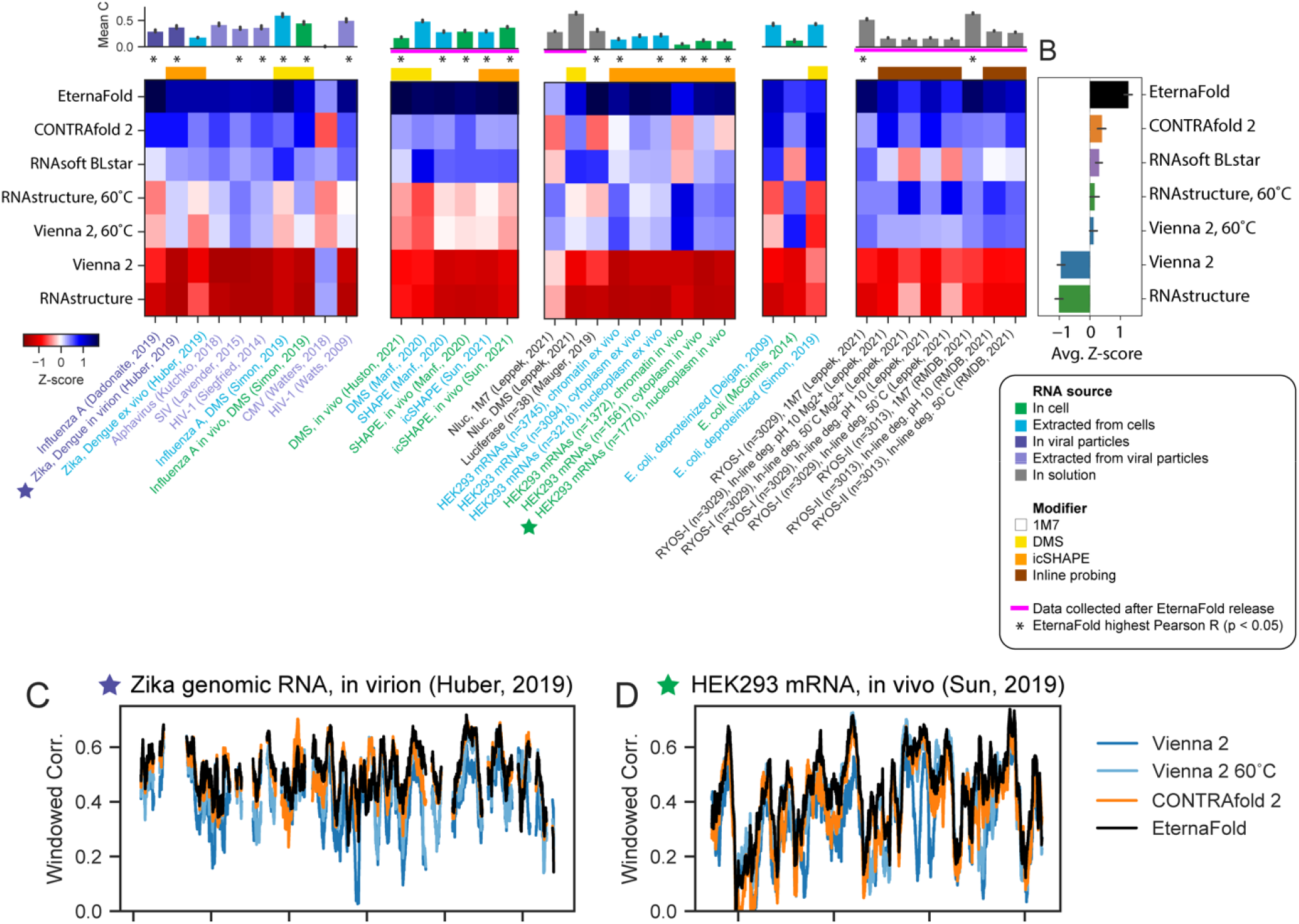
EternaFold improved prediction extends across diverse natural RNA contexts and experiments. A) Mean package correlation for top-performing representative packages selected to benchmark for 36 independent chemical mapping datasets from a variety of biological contexts and with other chemical modifiers, and is ranked highest in average Z-score (B). Calculation correlations on sequence windows indicates that EternaFold demonstrates uniformly higher correlation across sequence position for three representative datasets: (C) Zika genome probed in virion, (D) a human mRNA probed in vivo (ref. ^52^).

For 20/36 datasets across all categories, EternaFold exhibited the highest correlation coefficient (with p<0.05, determined by 95% overlapping CI, see Methods), and had the highest average Z-score (Figure 4B, Table 2). For the other 10 datasets, EternaFold was tied with other packages for having the highest correlation. EternaFold showed significant improvement (p<0.05) in datasets from varying sources including RNAs probed in cell (7/8 in cell datasets), extracted from cells (5/10), in virion (2/2), extracted from viral particles (3/5), and with other modifiers, including DMS (3/6) and icSHAPE (8/10). EternaFold was the top-scoring package (p<0.05) in 5 of the 6 datasets of novel SARS-CoV-2 data.

We were curious as to whether the differences in packages arose from consistent accuracy differences across all regions of these RNAs or from a net balance of increased and decreased accuracies at specific subregions of the RNAs, which might reflect particular motifs that are handled better or worse by the different packages. We calculated correlations along the length of example constructs -- the Zika ILM genome probed in virion^44^ (Figure 4C), a HEK293 nucleoplasm mRNA probed in vivo^52^ (Figure 4D) -- and observed that EternaFold correlations generally demonstrated a fixed improvement across compared packages across all regions, supporting a consistent accuracy improvement by this package.

We also tested the ability of EternaFold to predict the thermodynamics of binding of human Pumilio proteins 1 and 2 in a dataset of 1405 constructs^53^. EternaFold showed no significant increase or decrease in predictive ability (p>0.05) when compared to CONTRAfold or ViennaRNA 2 at 37°C (Figure S11, Table S10).

## Discussion

In this work, we have established EternaBench, benchmark datasets and analysis methods for evaluating package accuracy for two modeling tasks important in RNA structural characterization and design. These include 1) predicting unpaired probabilities, as measured through chemical mapping experiments, and 2) predicting relative stabilities of different conformational states, as exhibited in riboswitch systems. Unlike in single secondary structure prediction tasks, we demonstrate that both widely used and state-of-the-art machine-learning algorithms demonstrate a wide range in performance on these tasks. We averaged both rankings to acquire a final ranking of the tested external packages in Table 1.

We discovered that CONTRAfold 2, which inferred thermodynamic parameters by feature representation in datasets of natural RNA secondary structures, performed best in this ranking, and performed significantly better than Vienna RNAfold, NUPACK, and RNAstructure, packages with parameters derived from thermodynamic experiments^9^. The results were particularly notable since the probed RNA molecules were designed for two distinct tasks (chemical mapping and riboswitch binding affinities), with no relationship between these two sets of sequences and no relationship between the synthetic sequences and natural sequences. We further investigated if combining these tasks in a multitask-learning framework could improve performance. We found that models trained on four types of data – single structures, chemical mapping data, and riboswitch affinities for both protein and small molecules – showed improved performance in predictions for held-out subsets of EternaBench datasets as well as improvements in datasets involving virus RNA genomes and mRNAs collected by independent groups.

The improved performance of CONTRAfold and RNAsoft – two packages developed by maximum likelihood training approaches – was not obvious prospectively. Statistically-learned packages could incorporate bias towards common motifs in the RNA structures that they were trained on and might overstabilize motifs simply due to their increased frequency rather than actual thermodynamic stability. Indeed, methods developed with a variety of more recent methodological advances, including machine learning from chemical mapping datasets (CROSS), deep learning methods for secondary-structure prediction (SPOT-RNA), extended parameter sets (CONTRAfold-noncomplementary, CycleFold, MXfold), or accelerated folding packages (LearnToFold), demonstrated diminished performance in the EternaBench tasks (Figure S1). It was surprising that well-developed and more widely used packages like ViennaRNA and RNAstructure gave worse performance than CONTRAfold and RNAsoft across all tasks, but that predictions from ViennaRNA and RNAstructure at 60°C showed notable improvement over the default of 37°C. This observation might be rationalized by discrepancies in Mg^2+^-free solution conditions used to measure these packages’ thermodynamic parameters and the *in vitro* and *in vivo* conditions tested here.

We used the EternaBench datasets to train a thermodynamic model via multitask learning on secondary sturcture prediction, chemical mapping signal likelihood maximization, and minimizing error for riboswitch protein-binding prediction. The resulting model, termed EternaFold, performed best across 36 external datasets in 5 categories of natural and synthetic RNAs (Table 2) in a variety of cellular contexts, including RNAs probed in and extracted from cells and viral particles. Although many factors influence RNA structure in cells beyond thermodynamic base-pairing^54^, this demonstrates that existing natural RNA datasets are indeed capable of discriminating between ensemble-averaged base-pairing predictions, and that accurate prediction of chemical mapping signal presents an ensemble-aware target for RNA secondary structure algorithm improvement.

The improvements from multitask training in EternaFold indicated that the nearest-neighbor model encoded in CONTRAfold had sufficient representational capacity to gain improvement on the chemical mapping and riboswitch prediction tasks. Further improvements in modelling may arise from applying more sophisticated graph-^55^ and language-based^56^ architectures to predicting RNA thermodynamics. Further investigations may also be necessary to improve performance and aspects of the model that need to expanded, which may include noncanonical pairs^12^, more sophisticated treatment of junctions^57^, next-nearest-neighbor effects^14^, and chemically modified nucleotides^58^. Another area for improvement is the systematic evaluation of structure prediction methods that incorporate structure mapping data^8, 54, 59^, and will be enabled by high-throughput datasets that design orthogonal thermodynamic measurements in addition to chemical mapping. Taken together, the datasets presented here serve as an important starting point for evaluating and improving future RNA structure prediction algorithms.

## Acknowledgements

We thank members of the Das and Barna labs (Stanford University), C. Pop, and C.-S. Foo for useful discussions. We thank I. Jarmoskaite, V. V. Topkar, R. Wellington-Oguri, and J. Townley for helpful comments on the manuscript. Calculations and model training were performed on the Stanford Sherlock cluster. We acknowledge funding from the National Science Foundation (GRFP to H.K.W.S.), the National Institute of Health (R35 GM122579 to R.D.), and gifts to the Eterna OpenVaccine project from donors listed in Table S11.

## Contributions

H.K.W.S. and R.D. designed the EternaBench benchmark approach and EternaFold multitask training method. H.K.W.S. prepared the EternaBench datasets, performed analyses, and implemented and trained the EternaFold model. H.K.W.S. and R.D. wrote the manuscript. W.K. designed methods, acquired data for high-throughput chemical mapping experiments, and reviewed the manuscript. A.I.S. performed data analyses and visualizations. W.K., J.L., A.T., and R.D. designed and implemented the Eterna Cloud Lab initiative. Eterna participants created online design projects, provided RNA solutions and reviewed the manuscript (see Supplemental Table 1).

## Methods

The algorithms evaluated in this work model secondary structure in the following manner. Given a model *Θ*, which is comprised of a set of structural features {*θ*}, the partition function of an RNA sequence *x* is computed as

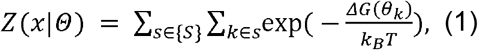

where *ΔG*(*θ*_*k*_) is the free energy contribution of structural feature *k, k*_*B*_ is Boltzmann’s constant, and *T* is temperature. Z represents a sum over the set of all possible structures {*S*} ^60^. From this expression, the probability of any particular structure *s* is defined as

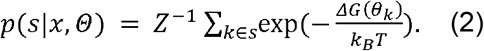

### Chemical mapping prediction theoretical basis

Structure prediction algorithms are able to estimate the ensemble-averaged probability that a nucleotide is paired or unpaired. Let *p*(*i*: *j*|*x, Θ*) be the probability of bases *i* and *j* being paired, given sequence *x* and model *Θ*. For simplifying notation, we continue with implicit *x* and Θ, i.e. *p*(*i*: *j*|*x*, Θ) = *p*(*i*: *j*). This is computed as

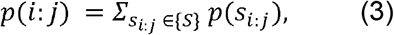

where *s*_*i*:*j*_ denotes a structure containing the base pair i:j, and {S} is the full set of possible structures. These posterior probabilities are analytically calculated by all the algorithms tested here. The probability of any single base being unpaired can be computed as

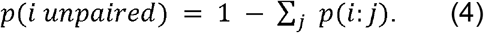

The relationship between the probability of a nucleotide being unpaired and its experimentally-measured reactivity has served as a locus for many efforts for improving structure prediction of RNA constructs incorporating chemical mapping data from those constructs, and several functional forms have been used to describe the relationship between unpaired probability and chemical mapping reactivity^27, 61, 62^. In this work, we use the linear Pearson correlation coefficient between unpaired probability and experimentally-measured reactivity as a measure of model quality. In the following, we describe the simple model under which this linear assumption holds. We write the probability nucleotide *i* is modified at time *t as*

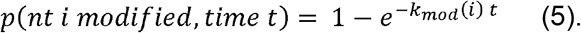

The measured chemical modification signal is an ensemble population average, where the time exposure of the ensemble to the modifier has been limited to aim to achieve “single-hit kinetics” with single-hit frequency, so that the degree of modification in experiment is proportional to the rate of modification^63^. In other words, because *k*_*mod*_(*i*) *t* ≪ 1, we can approximate

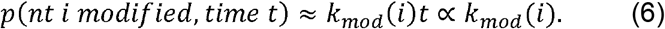

If we assume that the timescale of chemical modification is much slower than the timescale of fluctuation between structural ensemble states, then we may write the overall modification rate for each nucleotide *i* as averaged over the equilibrated structure ensemble of the RNA,

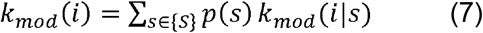

If we consider a simplest two-state model for each nucleotide, with modification rate *k*_*pr*_ if paired and a rate *k*_*unp*_ if unpaired, then this reduces to

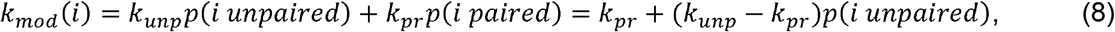

which demonstrates that under this simple model, the modification rate is linear with respect to p(unpaired). The model above is limited in its assumption of two states, and does not account for reactivity effects caused by sequence and local environment. A Spearman rank correlation (Figure S6), which will be more dominated by relative rankings, results in a similar overall ranking.

### Chemical mapping data

Chemical mapping data for the Eterna Cloud Lab experiments were downloaded from RMDB^**28**^ and processed with RDATKit (https://ribokit.github.io/RDATKit/). The RNA was probed with the MAP-seq protocol with a co-loaded standard molecule (P4-P6-2HP RNA) to enable normalization, as described in ref. ^64^; measurements were carried out at ambient temperatures (24 °C) with 10 mM MgCl_2_, 50 mM Na-HEPES, pH 8.0.

Within each chemical mapping dataset, CD-HIT-EST^**65**^ was used to filter sequences with greater than 80% redundancy (excluding a shared 3’ primer binding site). From each sequence cluster identified, the sequence with the highest signal-to-noise ratio from chemical mapping experiments was selected as the representative sequence. These datasets ranged in size from 605 (Round 15) to 3378 constructs (Round 23), with a median size of 1577; after filtering, they ranged from 101 (Round 12) to 1088 (Round 1), with a median size of 562 (Figure S3, Table S2). The filtered 24 datasets comprised 12,711 individual constructs, and distributions of GC content, average sequence length, and number of loops in the target structures were not significantly impacted (Figure S3).

Nucleotides with reactivities less than zero or greater than the 95th percentile of the dataset were removed from analysis. Cloud Lab Round 2 was filtered to exclude certain experiments that had FMN present, pertaining to Eterna Cloud Lab challenges to design riboswitches. Adenosine nucleotides preceded by 6 or more As were also removed due to evidence of anomalous transcription effects in such stretches ^**66**^, though this removal was not shown to alter package correlations to data (data not included). External chemical mapping datasets were obtained from the supplementary information from the papers and processed similarly (outliers, nucleotides in poly-A stretches removed).

### Analyzing package performance by Cloud Lab project

We wished to understand if factors such as target structure complexity, GC content, and sequence length influenced package predictions. We performed the same package ranking analysis, grouping constructs by their projects instead of by the 24 datasets. Because grouping constructs into projects sometimes resulted in a small number of nucleotides over which to calculate correlations, we omitted package predictions where the standard error of the calculated Pearson correlation was greater than 0.05. This resulted in a total of 612 project groupings remaining, names, and calculated metrics for which are contained in Table S4. We found weak correlation between the per-project Z-score of the top-performing package, CONTRAfold 2, and GC content (Spearman R = 0.15), sequence length (0.07), and total loops in the target structure (R=0.16). There were also weak correlations between the average Pearson Correlation for all packages and GC content (Spearman R = 0.10), sequence length (R=-0.24), and total target structure loops (R=-0.01) (Figure S7).

### Riboswitch activity prediction theoretical basis

A thermodynamic framework discussed in greater detail in ref. ^17^ allows us to relate the observed binding affinity of an output molecule to the relative populations of a riboswitch molecule in different states. In the absence of input ligand, we may relate the probability that a riboswitch adopts a structural feature that can bind its output, *p*(*out*), to an experimentally-measured binding affinity, 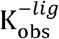, via the relative ratios of both values to those of a reference state:

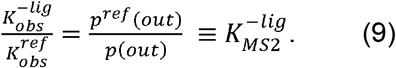

We selected the MS2 hairpin aptamer as a reference state whose probability of forming, *p*^*ref*^(*out*), can be estimated by the secondary structure algorithm. For each separate independent experimental dataset, 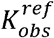 is estimated as the strongest affinity measured (Figure S12). We refer to the estimated ratio 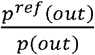 as 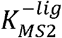 in the main text, as the equilibrium constant of forming the MS2 hairpin as normalized to the reference state.

Although there may be error introduced in which experimental point is selected to be 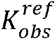, relative error should be constant when comparing packages on the same dataset. To compare packages, we report the correlation between 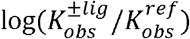 and 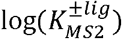, which excludes the effect of selection for 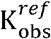.

In general, the probability of an RNA molecule forming any structure motif is computed as

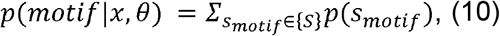

where *s*_*motif*_ denotes a structure containing that motif. Computing this probability requires a dynamic programming routine that is able to constrain the sampled structure space to only structures containing that motif to estimate a so-called “constrained partition function”. However, not all secondary structure algorithms have implemented constrained partition function estimation. Because the MS2 aptamer is a hairpin, we can approximate its probability of forming as the probability of forming the final base pair of the MS2 hairpin aptamer (colored pink in Figure 3A), an experimental observable that can be estimated by all the packages tested here. Thus, our prediction of interest is

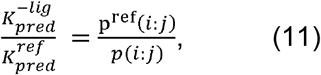

where *i* and *j* are the nucleotides forming the terminal base pair in the MS2 aptamer stem. The value *p*^*ref*^(*i*: *j*) is accordingly computed as the probability of closing the base pair in the reference sequence. We confirmed that calculations using eqn. 9 and eqn. 11 agree for Vienna, RNAstructure, and CONTRAfold packages.

### Predicting protein-binding affinities with input ligand bound

The estimation of 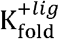 follows similarly to above but accounts for increased thermodynamic weights for states that correctly display the aptamer of the input small molecule ligand. Therefore, it cannot be estimated via the simplified single base pair calculation and must make use of constrained partition functions (eq. 10).

Analogously to eq. 9, we define 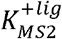 as

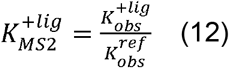

Which is calculated as

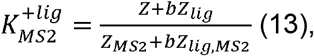

where 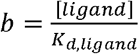, the Boltzmann weight of binding the ligand when the bulk concentration of the ligand is [ligand]. Values used for calculating *b* are in Table S12. Representative predictions of 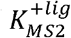 vs. experimental 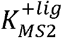 values are in Figure S13.

### Riboswitch data

Riboswitch data were downloaded from supplementary materials from refs. ^33^ (unpublished) and ^34^. In brief, measurements were carried out at 37 °C in 100 mM Tris-HCl, pH 7.5, 80 mM KCl, 4 mM MgCl_2_, 0.1 mg/mL BSA, 1 mM DTT, 0.01 mg/mL yeast tRNA, 0.01% Tween-20, and varying concentrations of small molecule ligand (FMN, tryptophan, theophylline) and MS2 coat protein. Datasets were filtered to only include constructs with sequences that included the canonical MS2 and small molecule aptamers, and filtered using CD-HIT-EST^65^ to remove sequence redundancy over 80%. Scripts to replicate data processing from refs. ^33^ and ^34^ are included in the EternaBench software.

For all constructs as well as the reference MS2 hairpin construct, we performed 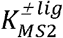 estimations including a flanking hairpin included in the Illumina array experiments, described in ref. ^33^ (unpublished). In example, the full reference MS2 hairpin construct, as well as the constraint used for estimating 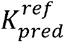 with constrained-partition-function-based estimation, is reproduced below. The MS2 hairpin construct is underlined and the nucleotides in the base used for base-pair-based prediction are bolded.

~~~
GGGUAUGUCGCAGAA**<u>A</u>**<u>CAUGAGGAUCACCCAUG</u>**<u>U</u>**AACUGCGACAUACCC
……………(((((x((xxxx)))))))……………
~~~

The riboswitches in EternaBench-Switch are controlled by the small molecules FMN, Tryptophan, or Theophylline. Motifs, concentrations, and intrinsic *K*_*d*_values used for 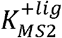 prediction, taken from refs. ^33^ and ^34^, are provided in Table S12.

### EternaFold Multi-task learning

The CONTRAfold^11^ loss function optimizes the conditional log-likelihood of ground-truth structure *s*^(*i*)^ given sequence *x*^(*i*)^ over dataset D:

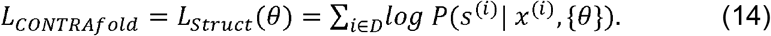

In CONTRAfold-SE^36^, the authors include a term to also use chemical mapping data to optimize structure prediction by maximizing the likelihood of observing the included chemical mapping dataset. The loss function then becomes

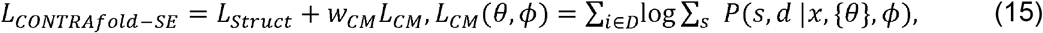

where *d* are the chemical mapping datapoints from construct *x*. CONTRAfold-SE fits reactivity signals to gamma distributions for each nucleotide type (A,C,G,U) and whether the base is paired or unpaired, parameters for which are represented by *ϕ*.

We further included a term to minimize the mean squared error of predicted 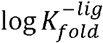 and 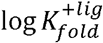:

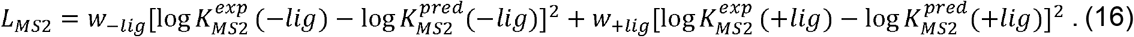

The full loss function for EternaFold is thus written as

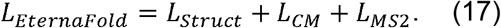

The hyperparameters *w*_*CM*_, *w*_−*lig*_, *w*_+*lig*_, corresponding to the relative weights placed on different data types, were selected through a grid search on a holdout set (data not shown). The final values used for training were *w*_*CM*_ = 0.5, *w*_−*lig*_ = 30, *w*_+*lig*_ = 30.

### Dataset selection for training EternaFold

#### Single-structure data

We used the S-Processed dataset^37^ train, holdout, and test sets used previously in training CONTRAfold 2 and RNAsoft^10^.

#### Cloud lab chemical mapping data

We used Rounds 3,4,5,7,10,11 as training and holdout data. This was to be consistent with the training data used for the CONTRAfold-SE work, and to reserve Rounds 0 and 1 as test rounds, given their large size and high signal-noise ratio. GC content, sequence length, total loops in the target structure, and signal/noise ratio were equivalent across train, holdout, and test rounds (Figure S14).

#### Riboswitch data

We partitioned the RiboLogic dataset into our training, holdout and test sets due to the high signal-noise ratio and diversity of structures, subdividing the riboswitches so that each split contained identical fractions of FMN-, Theophylline-, and Tryptophan-responsive riboswitches. This left the rest of the Eterna riboswitch rounds as test sets.

### Evaluating base-pair probabilities for external datasets

For comparing p(unpaired) calculations to natural RNAs, many of which are thousands of nucleotides long, we compared several practices for calculating, which includes predicting base-pair probabilities from overlapping windows, constraining the nucleotides under consideration using a beam search algorithm implemented in LinearPartition^67^, and conventional folding of the entire RNA. Windows of length 300, 600, 900, and 1200 with 25-nt overlap. Results from length 900 are shown in the main text, for which packages categories had the highest mean correlation, though results are similar for other window sizes (Figure S15). Figure S15 depicts correlation results from global folding for all package options tested.

### Error and Significance Estimation

We estimated confidence intervals on reported Pearson correlation values by bootstrapping the datapoints under consideration and reporting the 2.5^th^ and 97.5^th^ percentile over 1000 rounds of bootstrapping. Reported standard error values are estimated by calculating the standard deviation across bootstrapping rounds. We inferred significance in differences between package correlations by analyzing overlap between 95% confidence interval estimates^68, 69^. All code to reproduce significance analyses is included in the EternaBench repository.

### Code availability

The datasets used here for evaluation, as well as scripts and Python notebooks for reproducing the filtered datasets and the chemical mapping and riboswitch fold change calculations described here, are available at https://www.github.com/eternagame/EternaBench. The code for training EternaFold, as well as the training and test sets used, are available at https://eternagame.org/about/software as the package “EternaFold”. The EternaFold code is derived from the CONTRAfold-SE^36^ codebase, which is derived from the CONTRAfold^11^ codebase. A server to run EternaFold is being made available for noncommercial use.

### Package predictions

All base-pairing probability calculations and constrained partition function calculations were performed using standardized system calls through Python wrappers developed in Arnie (www.github.com/DasLab/arnie). Example command-line calls for each package option evaluated are provided in Table S1. Datasets were processed with Pandas (https://github.com/pandas-dev/pandas) and visualized with Seaborn (https://seaborn.pydata.org/).

## Supporting Information

**Figure S1:**
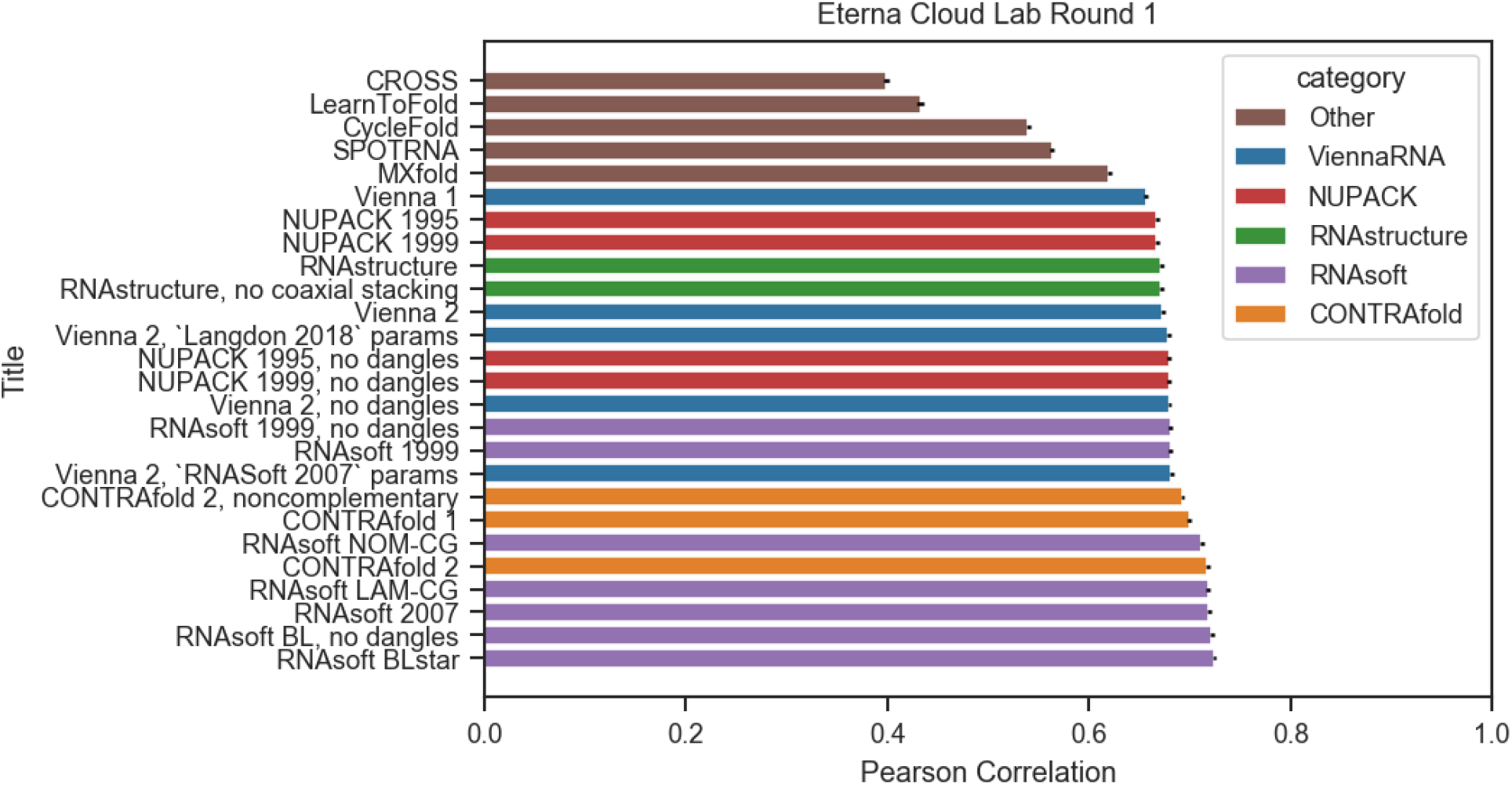
Correlation of all package options tested on Cloud Lab Round 69, which was also a holdout test set for EternaFold training studies.

**Figure S2:**
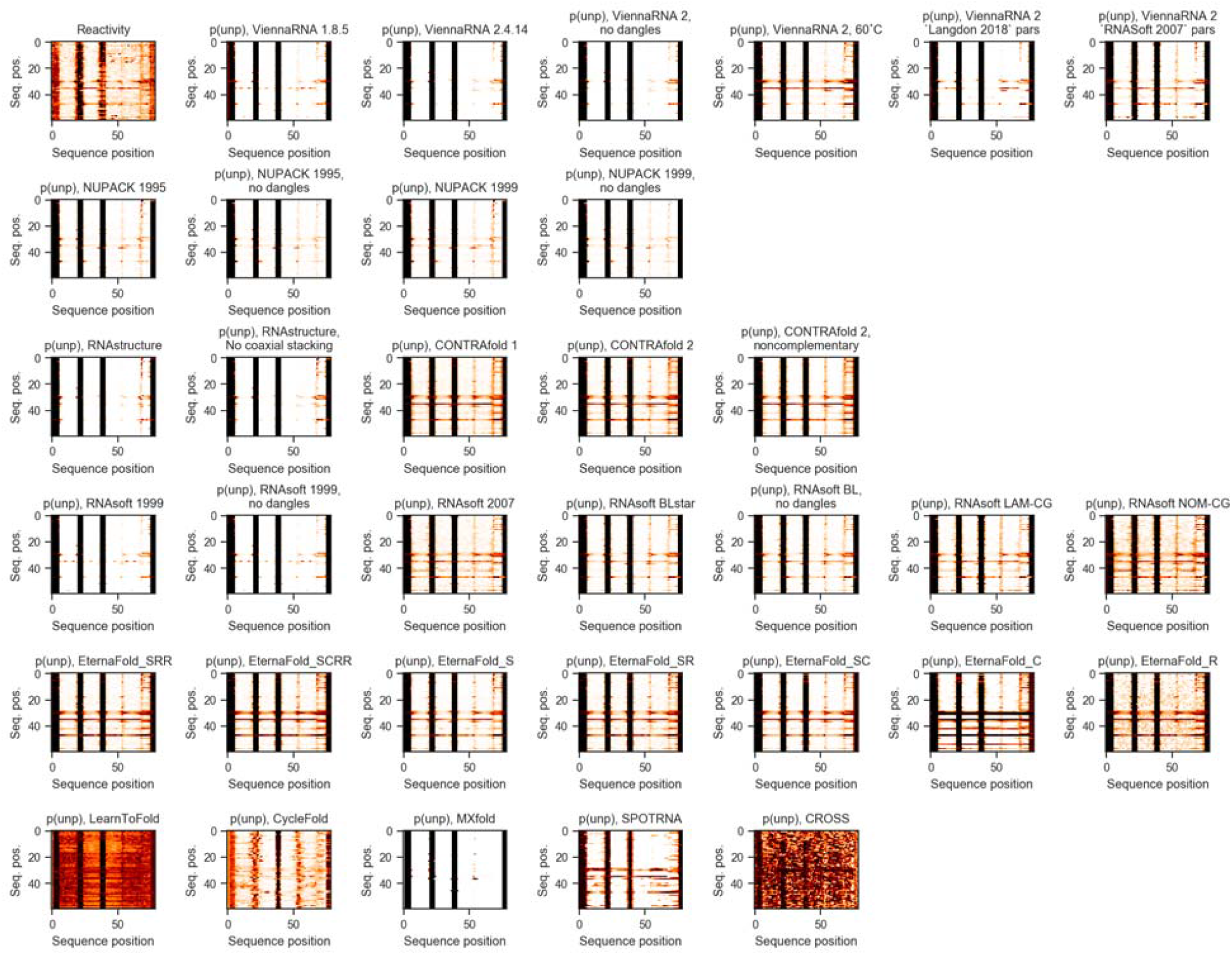
Example heatmaps of all package options tested, compared to heatmap of experimental reactivity data in panel 1.

**Figure S3.**
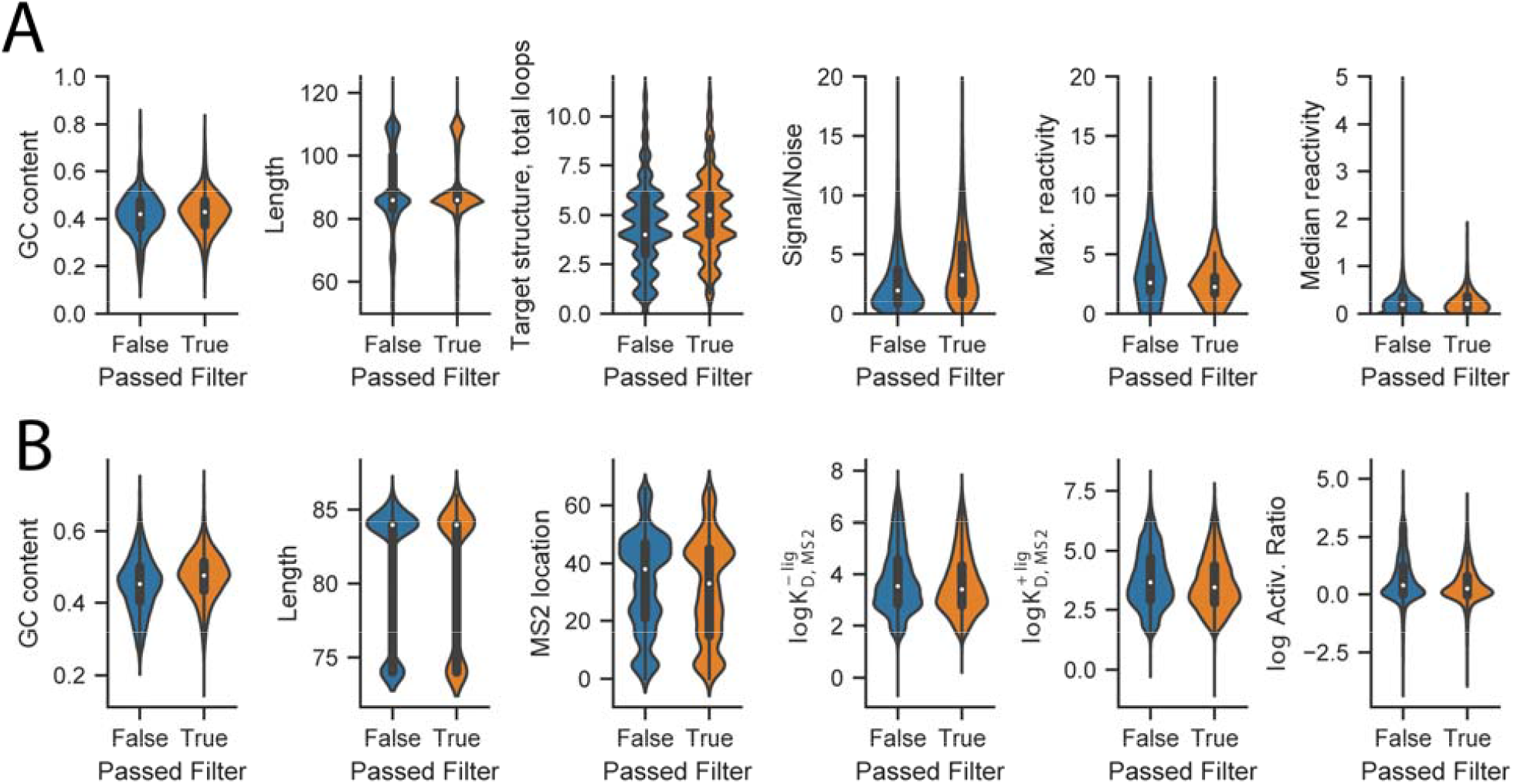
Summary statistics for EternaBench dataset CD-HIT filtering. A) Distributions of sequence properties for chemical mapping data, and B) riboswitch constructs were not affected by filtering.

**Figure S4:**
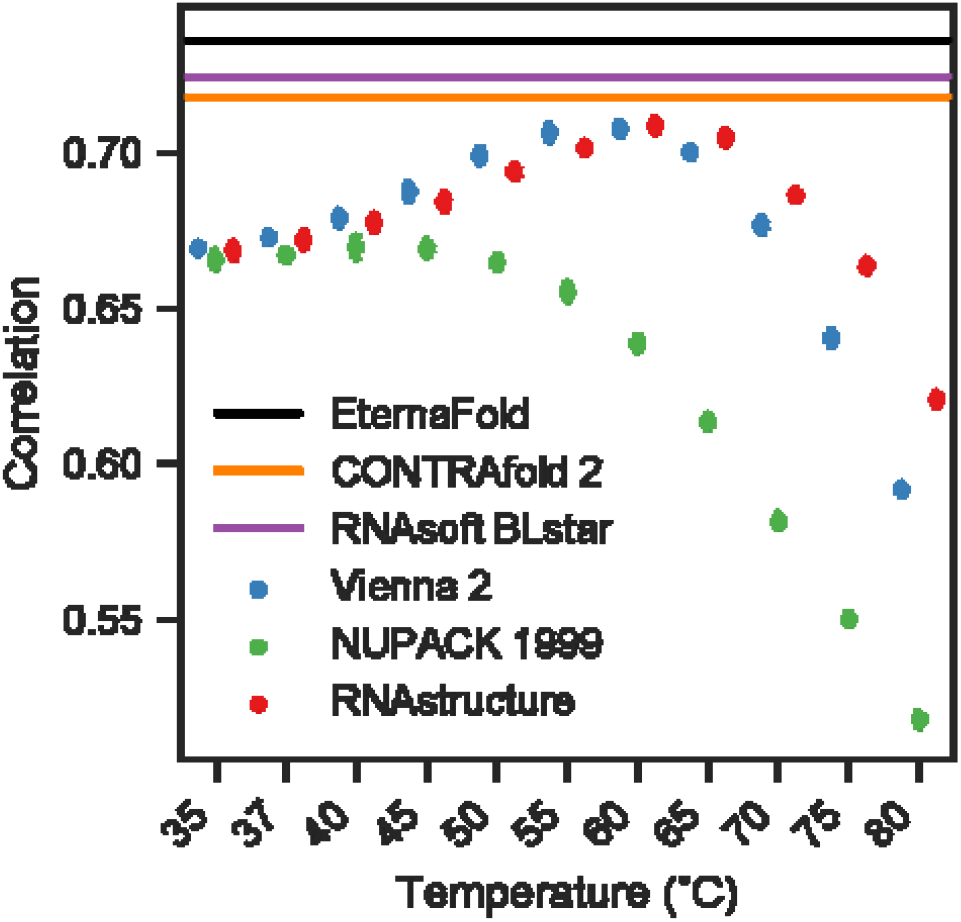
ViennaRNA 2, NUPACK 1999, and RNAstructure show maximum Pearson correlation to chemical mapping data at 60°C, 40°C, and 60°C respectively for Eterna Cloud Lab Round 1.

**Figure S5:**
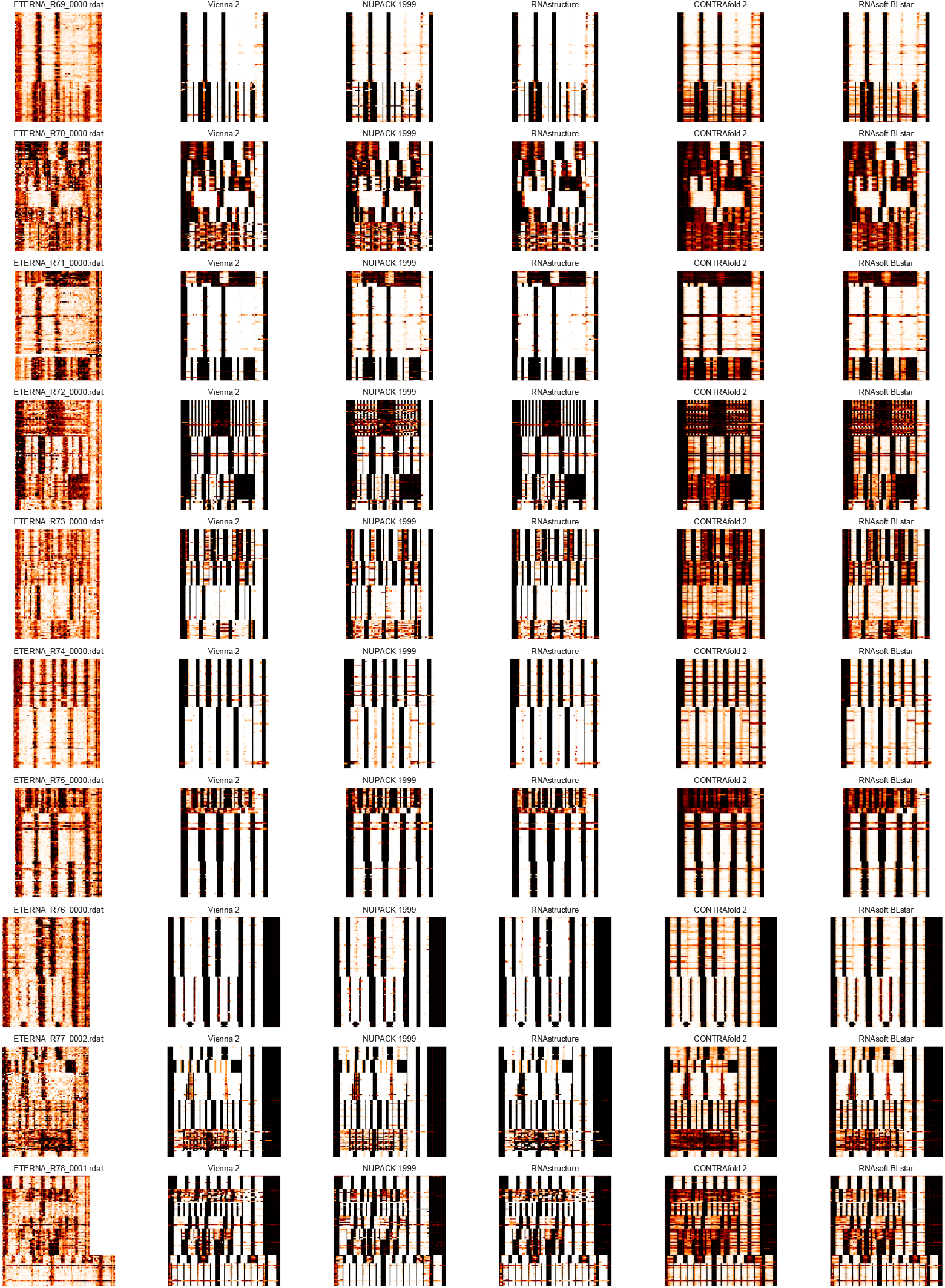
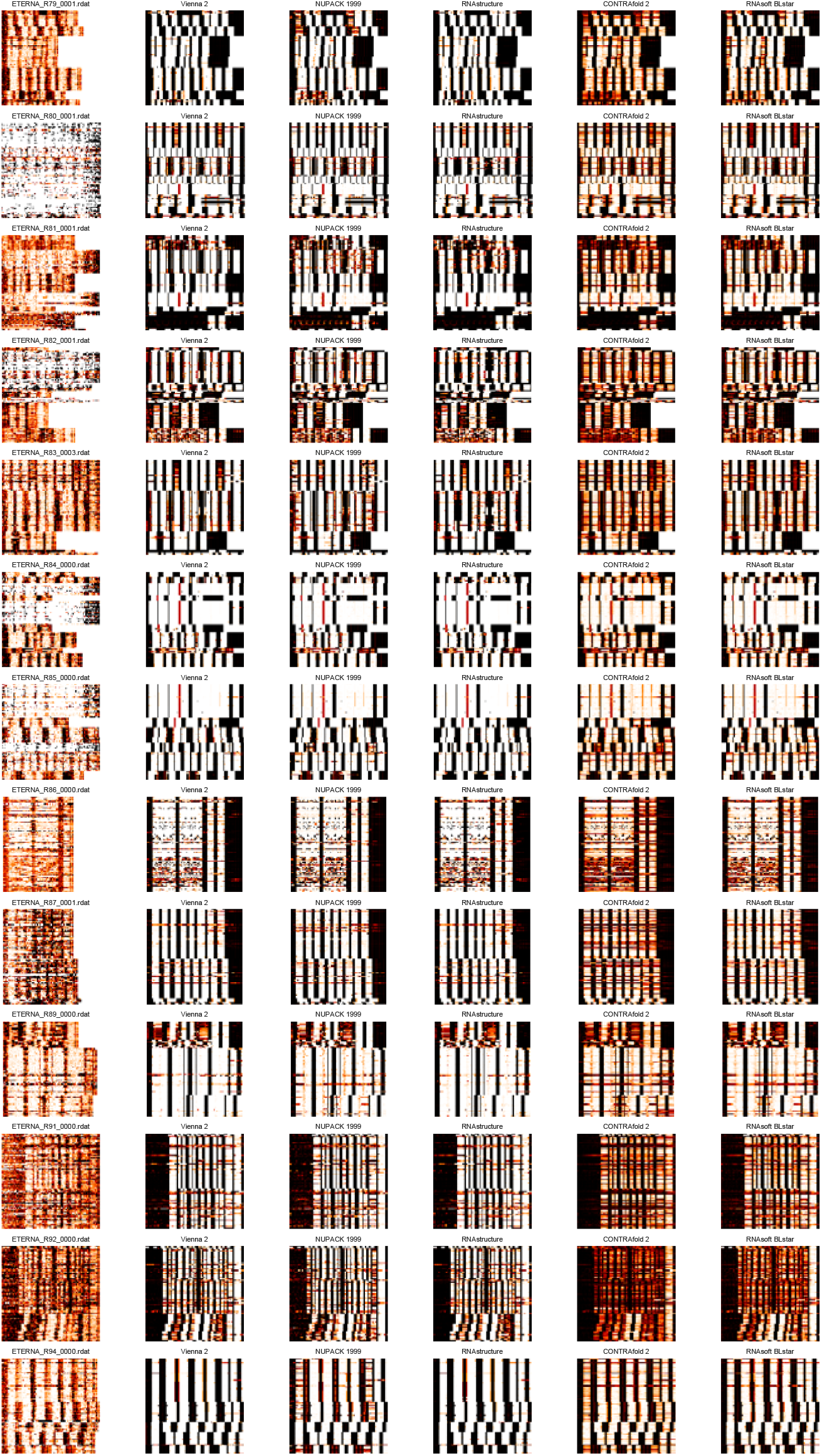
Example reactivity and p(unpaired) heatmaps from example packages for all 24 Cloud Lab rounds.

**Figure S6.**
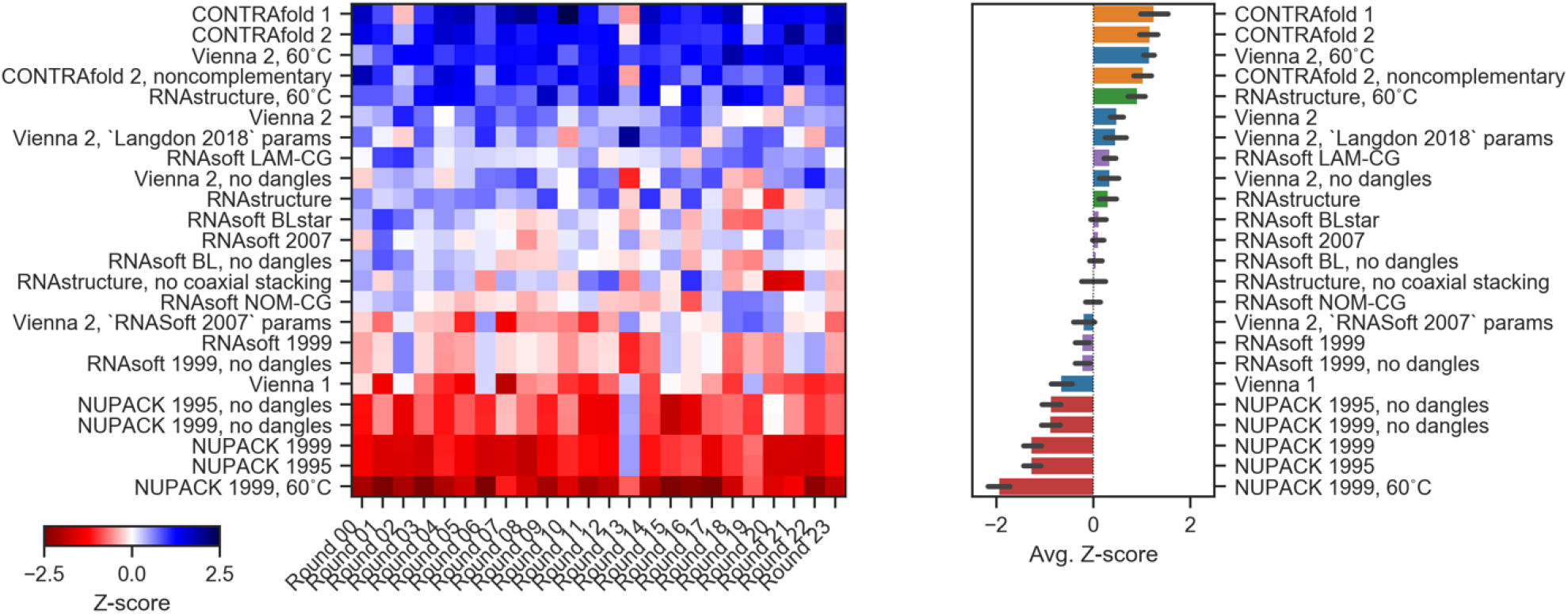
Ranking across Cloud lab dataset rounds using Spearman rank correlation (compare to Figure 1E,F).

**Figure S7:**
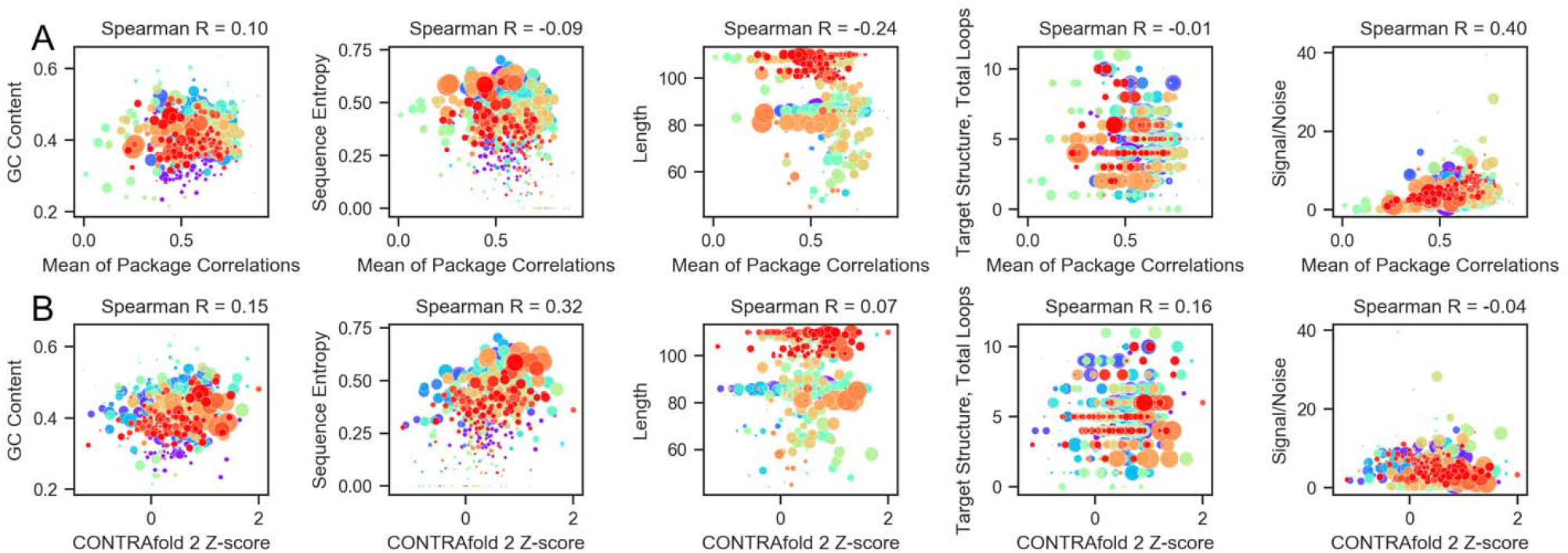
(A) Mean Pearson correlations, calculated over each project (as opposed to each dataset), compared to sequence metrics of the Cloud Lab projects. The strongest correlation to mean correlation was Signal/Noise ratio. (B) Z-score of CONTRAfold-2, calculated over each project, compared to sequence metrics of the Cloud Lab projects.

**Figure S8a:**
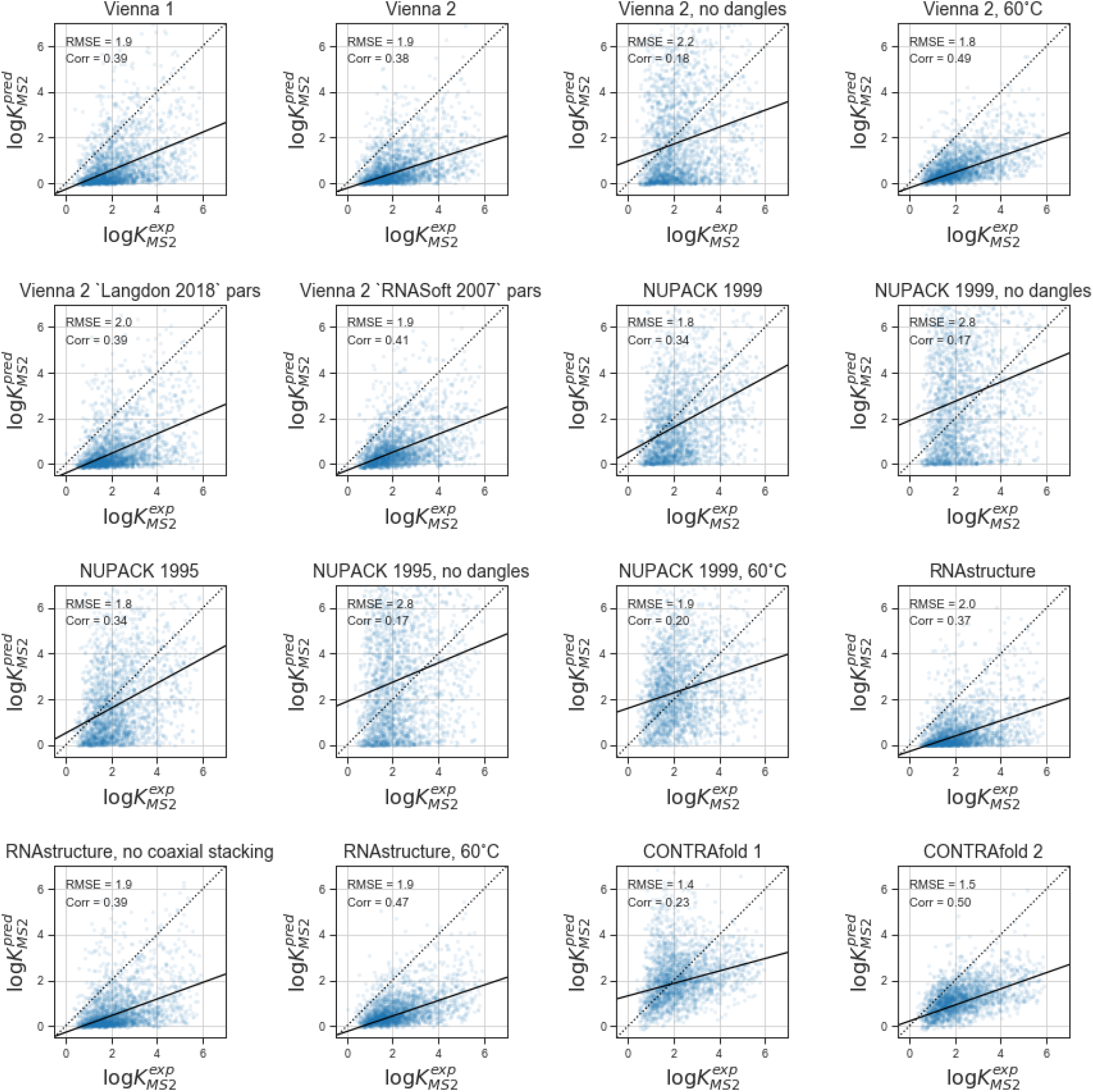
Scatterplots for all options tested for Ribologic dataset using closing-base-pair estimation (page 1 of 2). Black solid line indicates line of best fit.

**Figure S8b:**
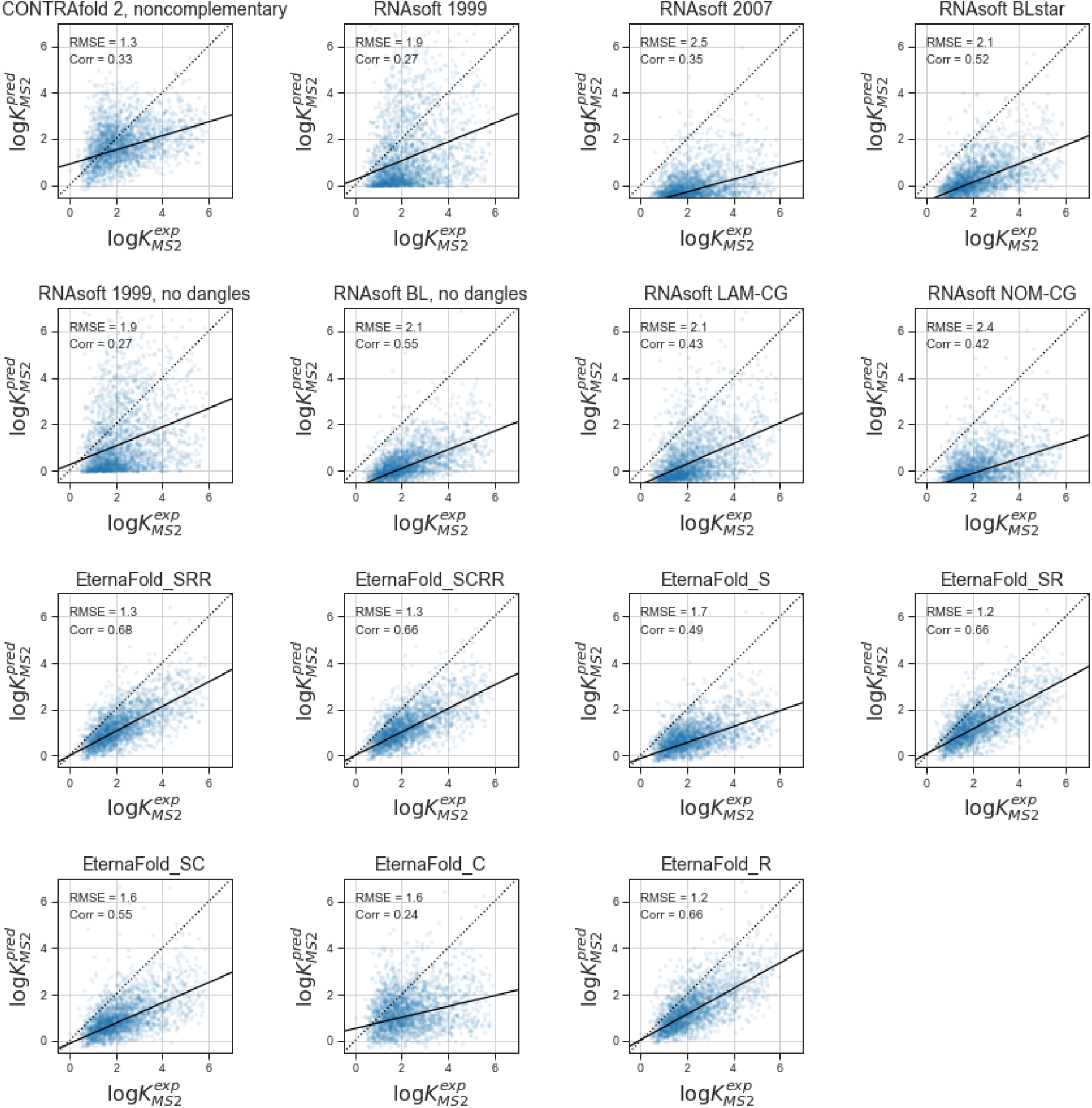
Scatterplots for all options tested for Ribologic dataset using closing-base-pair estimation (page 2 of 2). Black solid line indicates line of best fit.

**Figure S9a:**
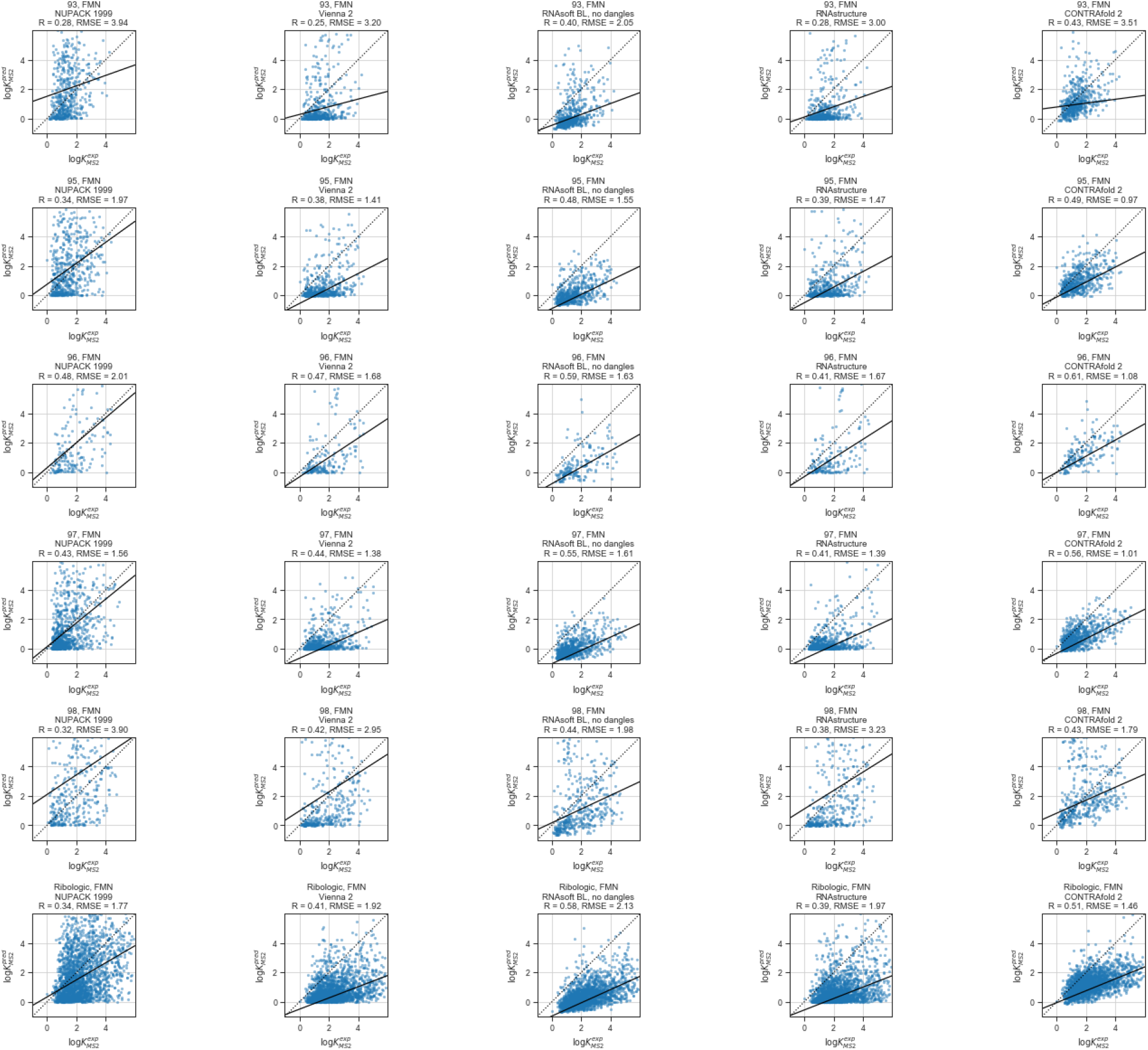
Scatterplots for representative packages on all riboswitch datasets (page 1 of 2). Black solid line indicates line of best fit.

**Figure S9b:**
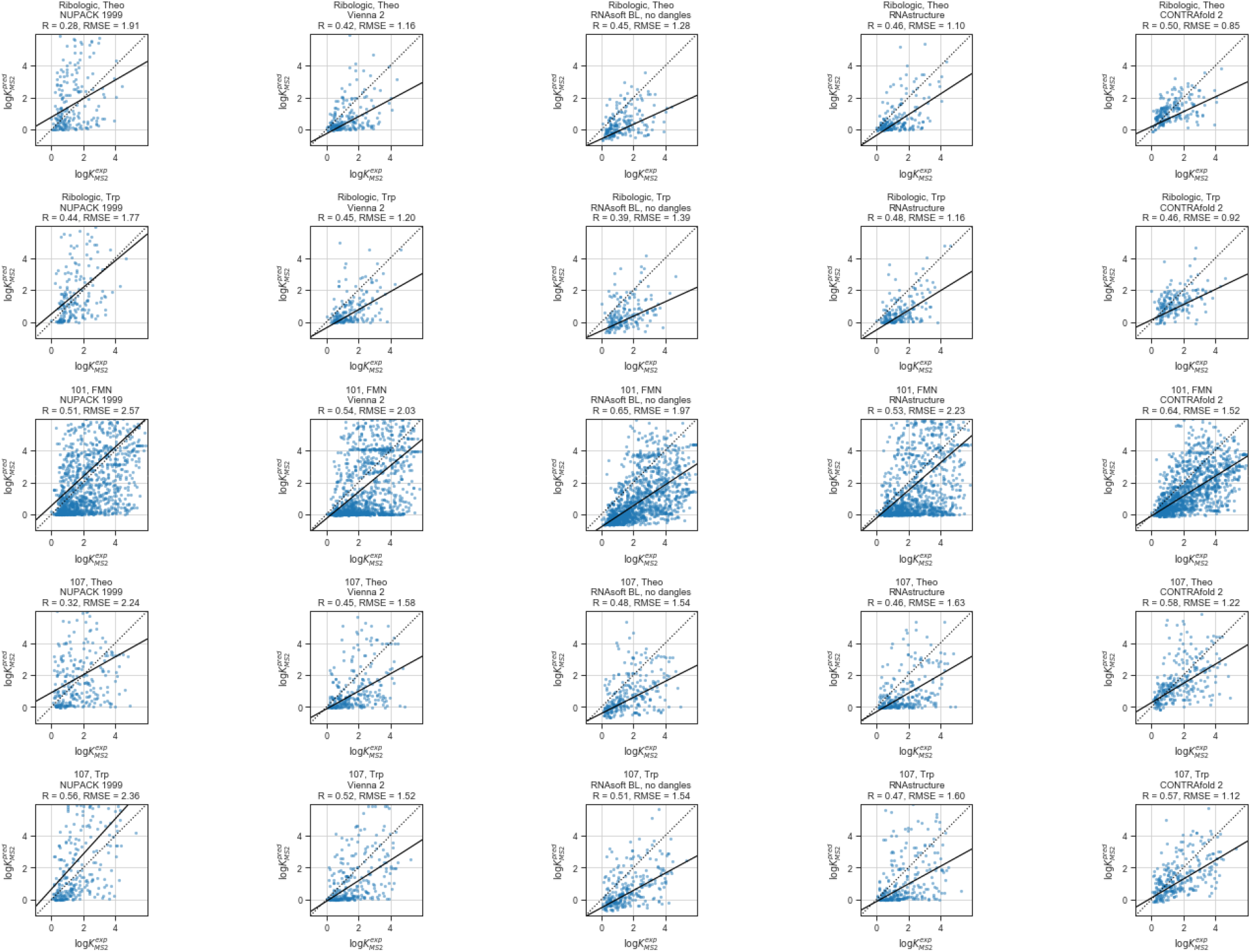
Scatterplots for representative packages on all riboswitch datasets (page 2 of 2). Black solid line indicates line of best fit.

**Figure S10.**
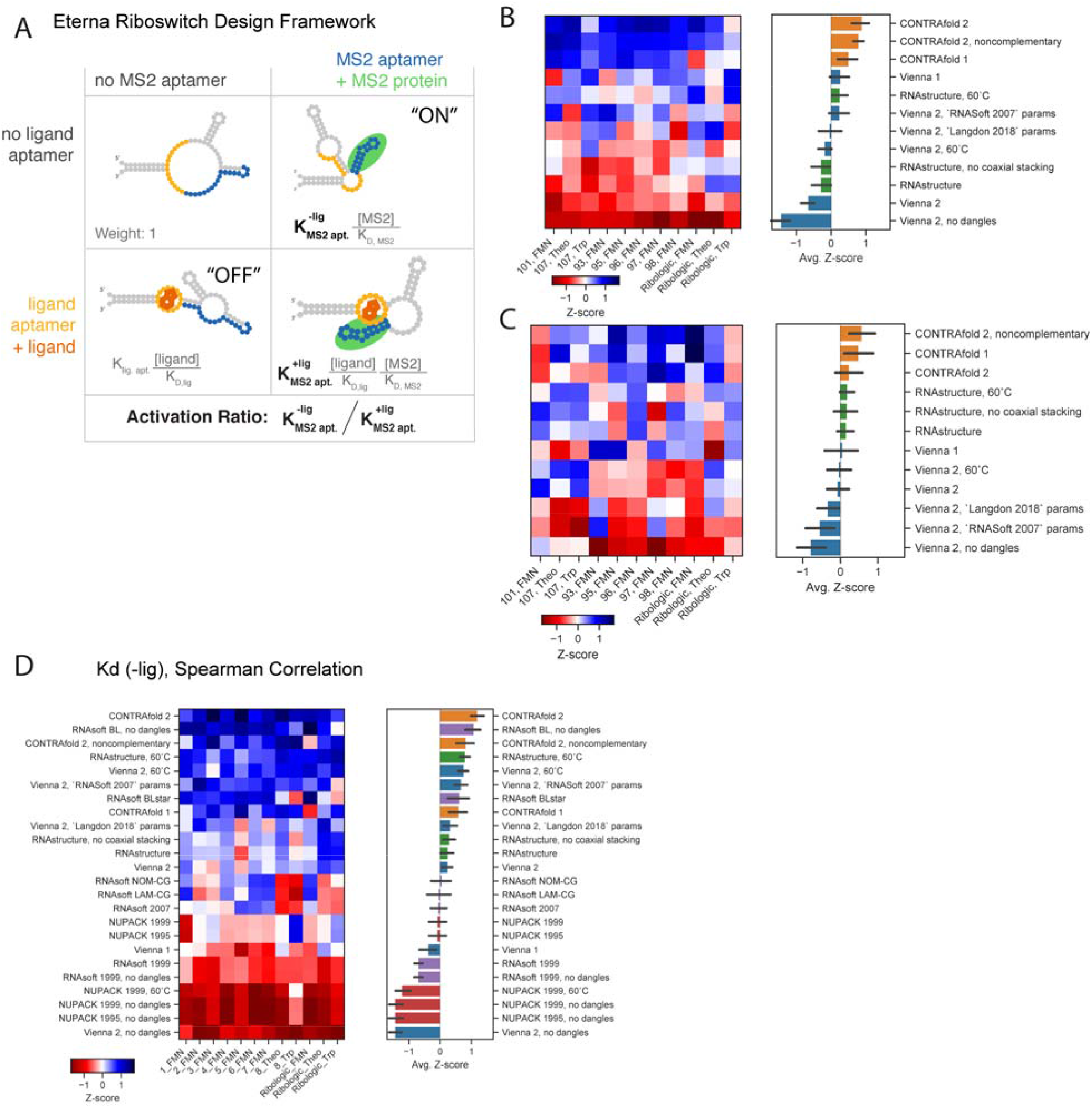
(A) Example set of states for a riboswitch that toggles binding of the fluorescent MS2 protein as an output, controlled by binding the small molecule FMN. The equilibrium constant for forming the MS2 aptamer in the absence of ligand, 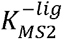, is estimated using the probability of forming the closing base pair for all packages. Evaulating the Pearson Correlation of package calculations for (B) 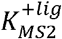 as well as (C) riboswitch Activation Ratio results in a similar ranking. (D) Overall ranking calculations using the calculated Spearman correlation (no linear assumption, compare to Figure 2B.)

**Figure S11:**
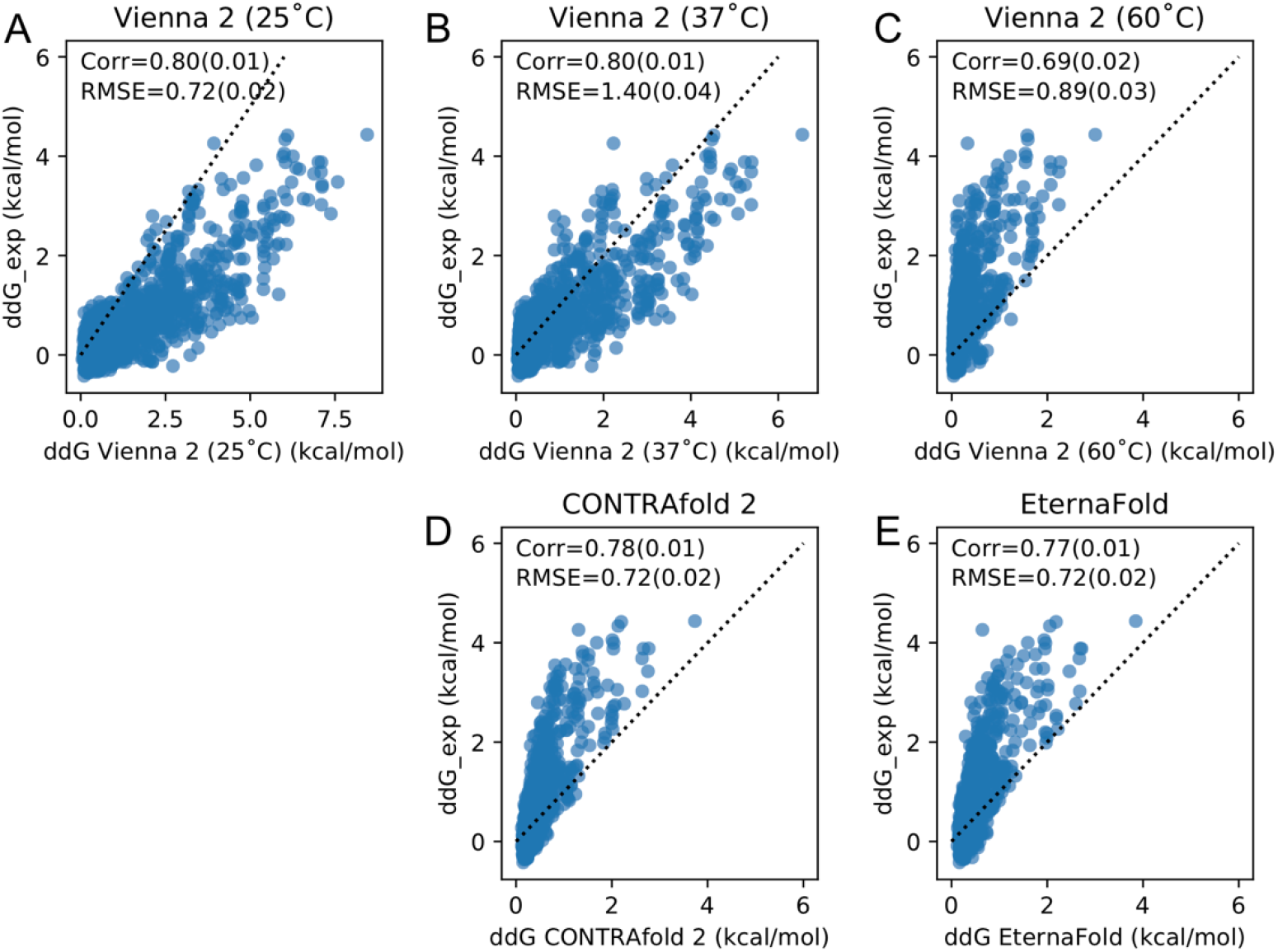
Comparing Vienna, CONTRAfold, and EternaFold predictions in predicting free energy of PUM binding. A) Replication of ddG_exp for both PUM WT and mutant binding from reference ^1^. The same calculation in Vienna 2 at 37°C shows lower RMSE (B), but higher RMSE at 60°C (C). CONTRAfold 2 shows no improvement over Vienna at 37°C (D), but EternaFold shows modest improvement over both (E).

**Figure S12:**
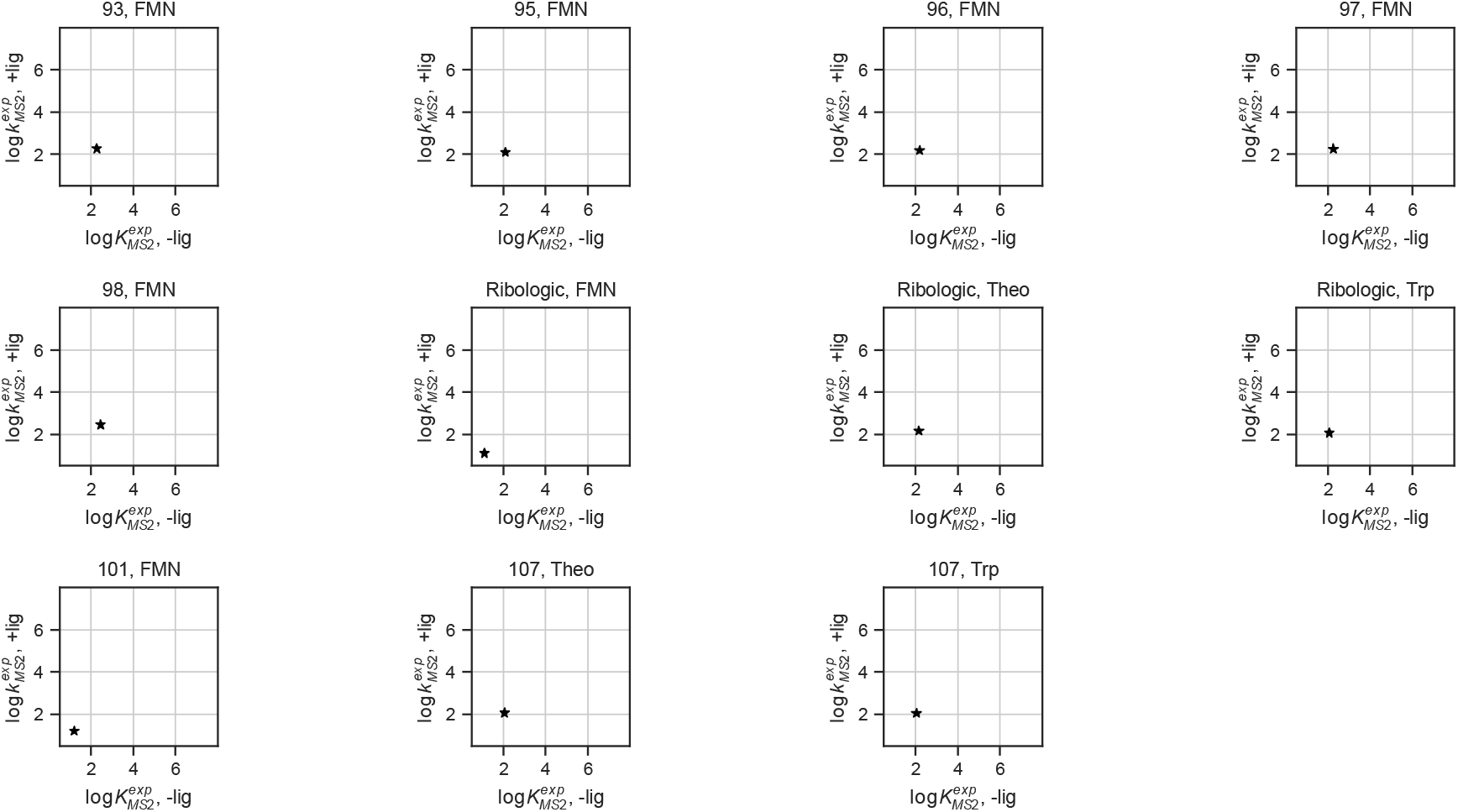
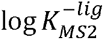 and 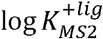 values of riboswitches included in filtered datasets. Black starred datapoint indicates reference value used for 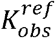.

**Figure S13:**
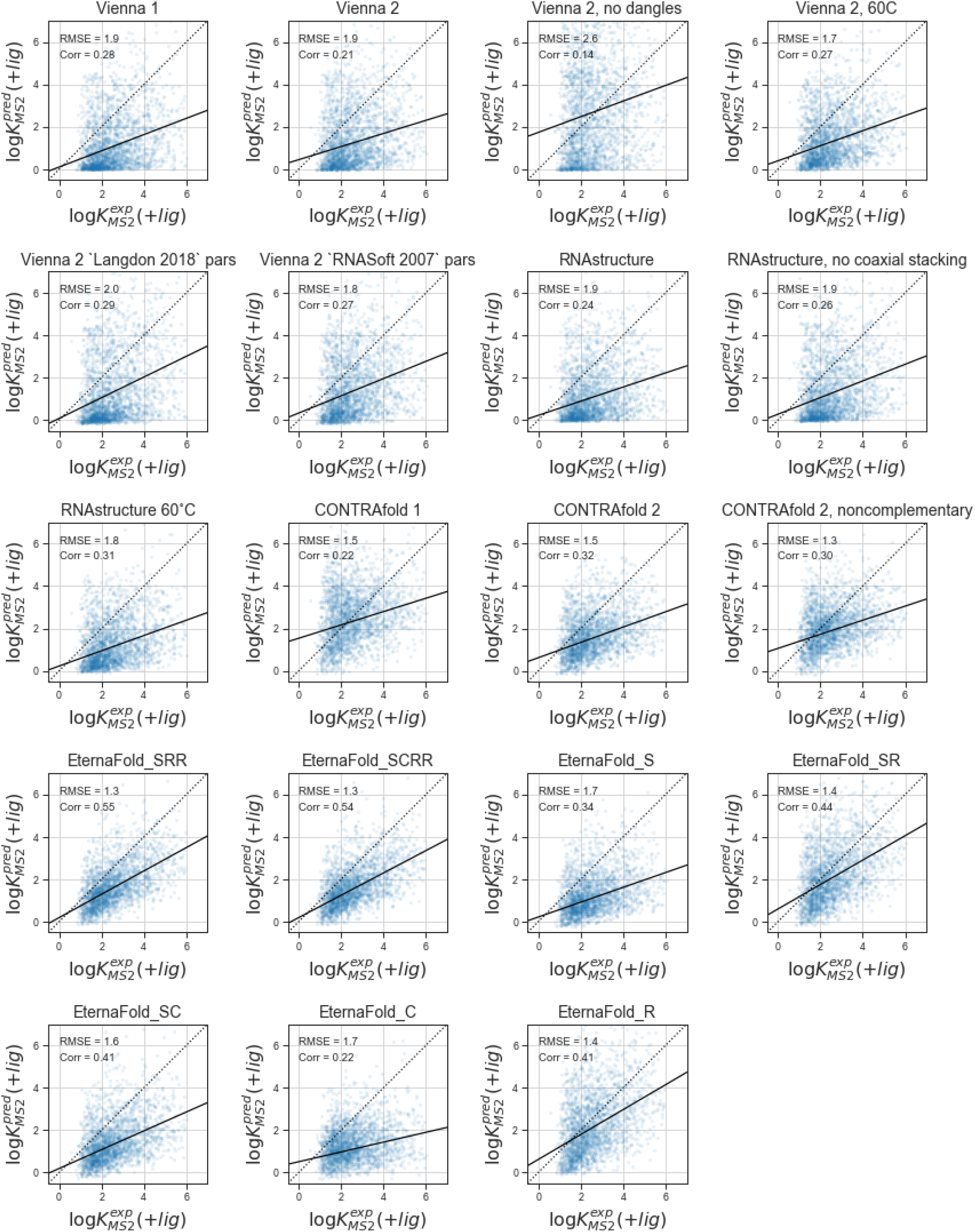
Estimates for the RiboLogic FMN dataset for in all package options able to make estimates with constrained partition functions.

**Figure S14.**
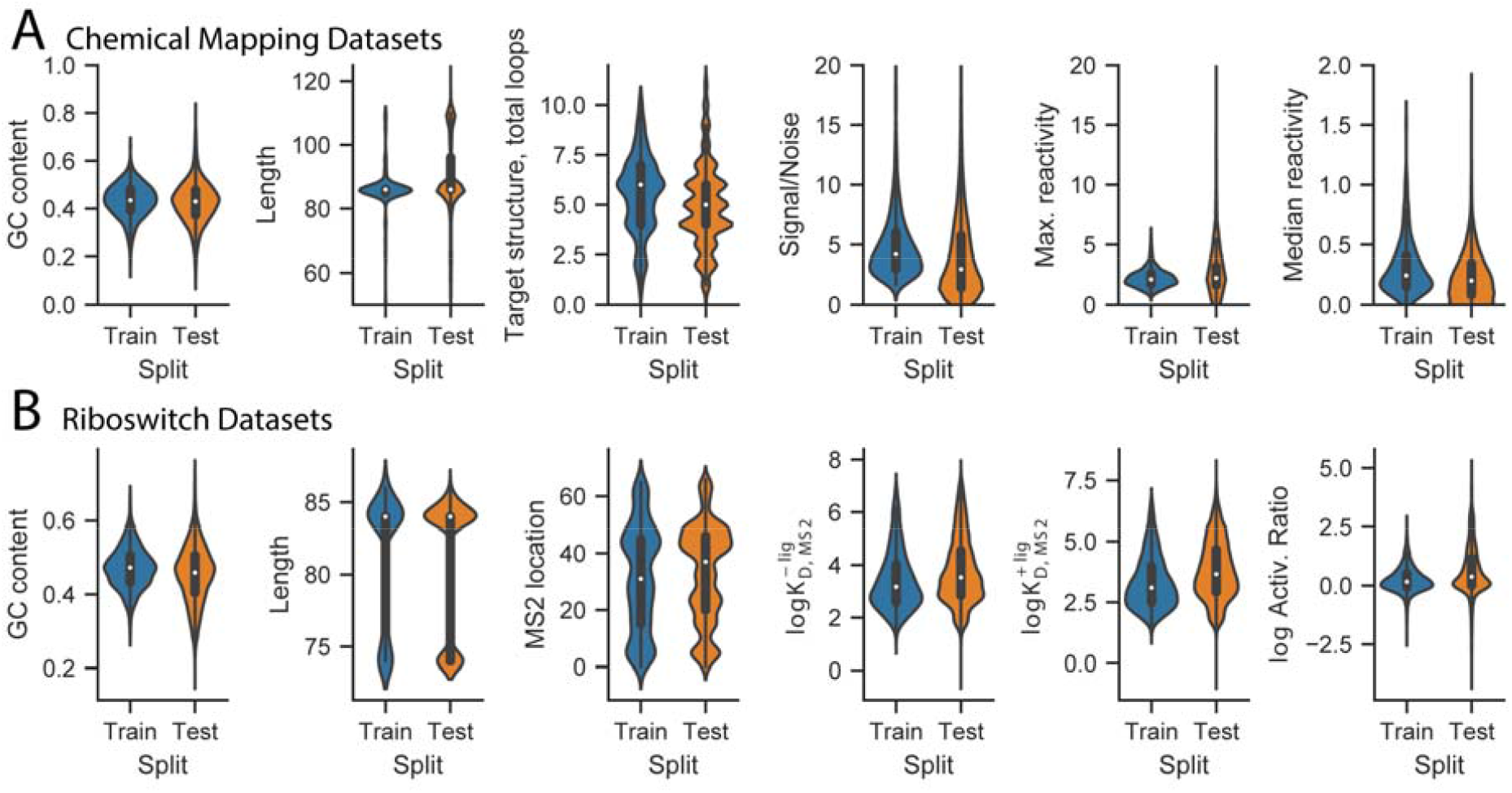
Dataset statistics of EternaBench train and test splits for (A) Chemical Mapping and (B) Riboswitch data.

**Figure S15.**
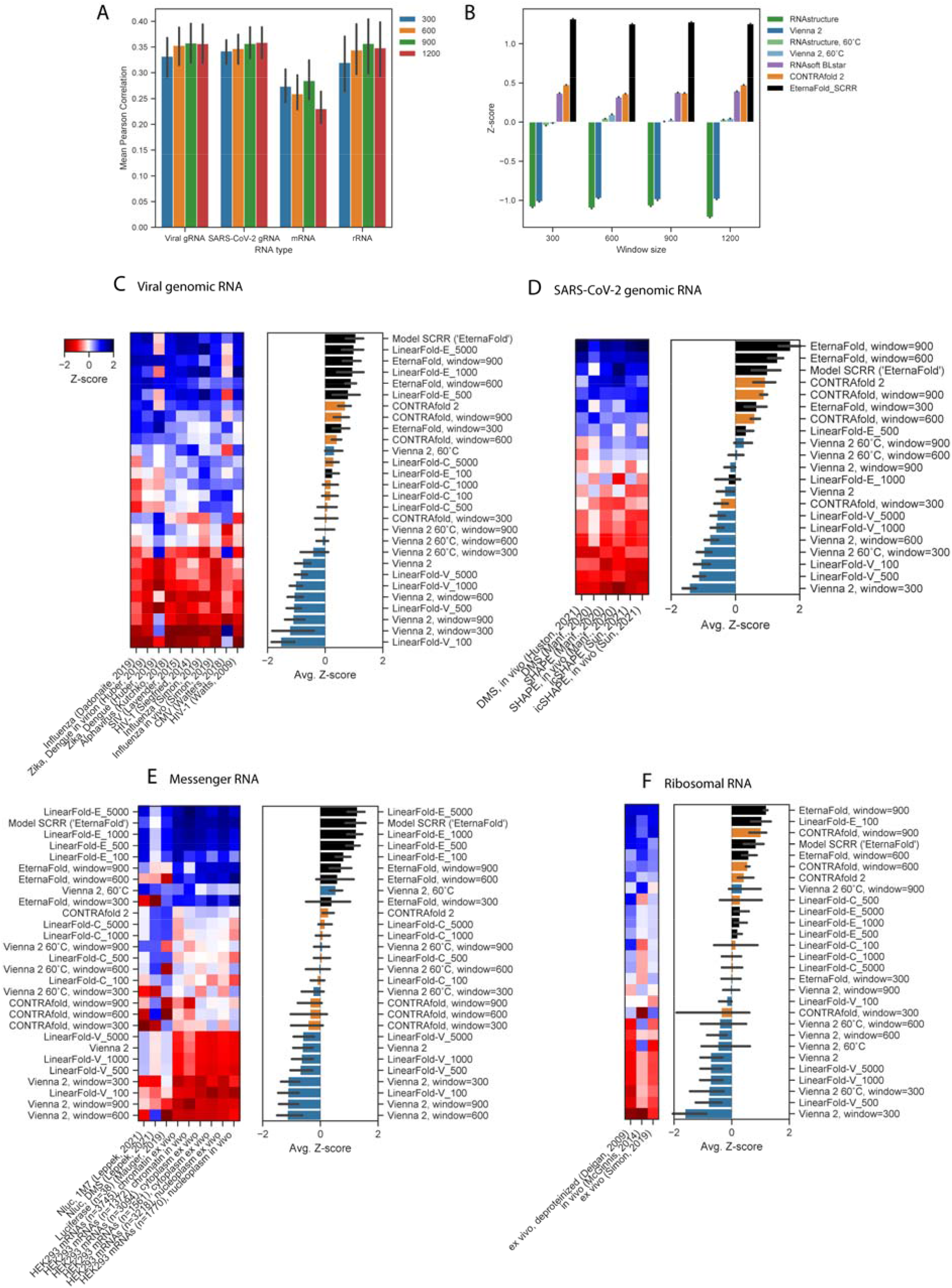
External data were folded using windows of 300, 600, 900, and 1200. A) Window size does not significantly change the correlation observed except for the mRNA datasets, which had the highest correlation at a window size of 900. B) Package ranking is consistent for different window sizes. Viral genomic RNA (C), SARS-CoV-2 genomic RNA (D), mRNA (E), rRNA (F), all options tested for global thermodynamic folding vs. LinearPartition with varying beam sizes and windowed folding. Fewer package options were tested for the SARS-CoV-2 genome due to overflow errors.

